# TurboID proximity labeling reveals protein interaction networks for germline Argonautes

**DOI:** 10.64898/2026.09.21.753052

**Authors:** Sebastian Fuentes-Otalora, Samantha Del Borrello, Ismail I. Irshaid, Isaac Martinez Ugalde, Gwynneth McLeod, Veronica Venditti, Cassandra J. Wong, Anne-Claude Gringas, Julie M. Claycomb

## Abstract

In the germline, small RNA-mediated gene regulatory pathways are associated with phase separated condensates known as germ granules across species. Germ granules can be challenging to purify based on their biophysical properties, therefore we used an *in vivo* proximity labeling approach, TurboID, to define the proximal interaction networks of the germ granule-localized Argonaute proteins: CSR-1, PRG-1/Piwi, PPW-2/WAGO-3, WAGO-1 and WAGO-4, and the nuclear-enriched Argonaute HRDE-1 as a comparator, in *C. elegans*. We found that although CSR-1 and WAGO-4 bind to the same set of 22G-small RNAs and target an overlapping set of germline genes, they possess distinct proximal interaction networks. Displacement of WAGO-4 from the germ granules highlights distinct interactors involved in chromosome biology and points to a potential nuclear role for WAGO-4. PRG-1/Piwi, WAGO-1, and PPW-2/WAGO-3, act in a common pathway and share an interaction network, while HRDE-1 displays a distinct interaction network that highlights its potential to influence splicing, rather than its widely accepted role in histone modification. Performing TurboID on other sRNA pathway factors, including the RdRP RRF-1 led us to uncover a potential nuclear role for this protein, and overlapping our TurboID data with published proximity labeling data enabled us to identify a core set of eleven mostly uncharacterized proteins that play roles in fertility, germ granule organization, and transgenerational epigenetic inheritance. One of these factors, D2005.4, appears to be a heme-binding germ granule-localized protein with a Tetratricopeptide Repeat Domain that interacts with all germ granule AGOs and may play scaffolding or enzymatic roles in germ granules. Collectively, our work highlights the utility of proximity labelling to uncover insights into AGO and sRNA pathway factor function and identify new factors that could be of importance in germ cell biology.

## INTRODUCTION

Maintaining optimal fertility is an intricate and complex process that ensures the survival of sexually reproducing species. Small RNA (sRNA)-mediated gene regulatory pathways contribute to the generation of functional gametes across plants and animals^1^. These pathways play multifaceted roles in suppressing transposable element mobilization, maintaining genome integrity, performing surveillance of potentially harmful foreign nucleic acids, and tuning the levels of germline transcripts^2,3^. In these pathways, sRNAs associate with Argonaute (AGO) proteins and guide AGOs to target transcripts by sequence complementarity, which leads to target regulation through a variety of mechanisms such as mRNA degradation, chromatin modification, and translational repression. The mode of action for sRNA pathways depends on factors such as the degree of base pairing between the sRNA and target, as well as features intrinsic to the AGO, including subcellular localization and association with different cofactors^4^.

*Caenorhabditis elegans (C. elegans)* has been a longstanding model for understanding sRNA pathway functions. The worm possesses an expanded repertoire of 19 functional AGOs and four types of sRNAs, of which, 16 AGOs and all four types of sRNAs are expressed in the germline^3^. Notably, half of these germline AGOs stably associate with non-membrane-bound organelles known as germ granules^5^. Germ granules are present across animal species and are generally required for proper germ cell development and differentiation, leading to optimal fertility^6^. In *C. elegans*, specific germ granules also play a role in the transmission of epigenetic information from parent to progeny, called Transgenerational Epigenetic Inheritance or TEI^5^. Altogether, *C. elegans* germ granules encompass six different subcompartments: P granules, Z granules, D granules, E granules, Mutator foci, and SIMR foci^4^. P granules are associated with the majority of nuclear pore complexes and are commonly associated with the storage and routing of germline mRNAs to other granules^7^. Mutator and SIMR foci are implicated in the amplification of specific subsets of sRNAs, known as 22G-RNAs^8,9^. Z granules are important for the inheritance of epigenetic information over generations via TEI^10^. Finally, E granules and D granules are thought to play roles in the generation of 22G-RNAs and mRNA export, respectively^11–13^.

These subcompartments house varying numbers of previously identified proteins, however the exact number and identity of proteins in each granule remains unclear due to the fact that many components are shared by multiple granules, proteins may move between granules, and, importantly, that not all germ granule components have been identified. Nevertheless, germ granules concentrate a variety of sRNA pathway factors including RNA dependent RNA polymerases (RdRPs), helicases, RNA degradation machinery factors, RNA binding proteins, AGOs, and RNAs^5^. Importantly, germ granules are devoid of ribosomes, suggesting that they are hubs of mRNA storage and regulation^14–16^. In *C. elegans,* germ granule-localized AGOs include CSR-1, WAGO-4, WAGO-1, PRG-1, PPW-2, ALG-3, ALG-4, and ALG-5^3,5^. PPW-1 is also expressed in the germline, however its localization is predominantly cytoplasmic and it is not enriched in condensates^3^. While these AGOs share the common feature of association with germ granules, they differ in their functions, expression patterns, and precise germ granule localization.

PRG-1 is the sole PIWI protein in *C. elegans*, and associates with tens of thousands of piRNAs, also known as 21U-RNAs, to perform genome and transcriptome surveillance in the germline^17^. Activation of the piRNA pathway by foreign nucleic acid leads to the production of an abundant class of small RNAs that are antisense and fully complementary to their target transcripts, known as 22G-RNAs^18^. 22G-RNAs are synthesized by the RNA dependent RNA polymerases, RRF-1 and EGO-1^19,20^, and loaded into a class of worm-specific AGOs known as WAGOs, including WAGO-1, PPW-2/WAGO-3^21^, and HRDE-1^22^. In this pathway, WAGO-1 and PPW-2/WAGO-3 operate in the cytoplasm, while HRDE-1 transits to the nucleus where it silences gene expression co-transcriptionally. In contrast to the post-transcriptional and co-transcriptional silencing capacity of the PRG-1/WAGO pathway, the AGO CSR-1, which is essential for fertility, is thought to protect or license approximately five thousand critical germline transcripts via its 22G-RNAs^3,19–23^. Notably, about 80 % of WAGO-4 22G-RNAs and target transcripts overlap with CSR-1, while a smaller subset of WAGO-4 targets overlap with the PRG-1/WAGO pathway^3^. Overall, WAGO-4 is thought to silence its targets, as loss of *wago-4* leads to an increase in target mRNA steady state levels^24^. A recent study also demonstrated that failure to recruit WAGO-4 to germ granules led to a shift in its pool of associated sRNAs, in which 22G-RNAs specific to the WAGO pathway decreased and 22G-RNAs overlapping with CSR-1 increased in association with WAGO-4^25^. This is consistent with the notion that WAGO 22G-RNAs are produced and loaded into WAGOs in germ granules, while CSR-1 class 22G-RNAs are generated and loaded in the cytoplasm.

Our knowledge of which germ granules each AGO localizes to has improved with advanced imaging methods and an ever-increasing number of fluorescently tagged germ granule proteins. Work from our lab and others indicates that PRG-1, WAGO-1, PPW-2/WAGO-3, WAGO-4 and CSR-1 localize to both P and Z granules to varying extents^3,5^. Recent higher resolution studies examining the overlap between fluorescently tagged candidate proteins with a set of designated markers for each germ granule demonstrate that WAGO-1 is enriched in P granules, PRG-1 in Z granules, and CSR-1 in D granules^13^. While HRDE-1 predominantly resides in the nucleus, it has been shown that impairing the 22G-RNA loading of HRDE-1 results in the accumulation of HRDE-1 in SIMR foci, indicating that HRDE-1 sRNA loading occurs in granules and is required for its nuclear localization^26^. As highlighted by the examples above, AGOs have dynamic activity cycles that lead to their recruitment to different subcellular compartments and granules. Little is known about AGO dynamics in the germline so far, therefore, defining which proteins each AGO interacts with *in vivo* may help us better understand these dynamics and the functional “life cycle” of the germline AGOs.

Phase-separated condensates such as germ granules present challenges to purification via conventional biochemical approaches^27^. Further, while immunoprecipitation-mass spectrometry (IP-MS) has been a valuable tool in identifying potential interactors of germ granule proteins such as AGOs, this approach has several drawbacks, including its inefficiency in capturing weak or transient interactions and the potential to introduce artifacts during the lysis procedure, as well as a loss of spatial context^28^. Instead, newer *in vivo* proximity protein labelling methods, such as BioID and TurboID, have become powerful tools in mapping the interaction networks of subcellular compartments such as germ granules^29–32^. In TurboID, a mutant version of the *E. coli* biotin ligase BirA is fused to a protein of interest (“direct” TurboID) or to a nanobody that recognizes a tagged protein of interest (“indirect” TurboID) and creates a reactive biotin species that labels exposed primary amines of proteins within approximately 10 nm^33^.

Here, we use indirect TurboID to define the proximal protein interaction networks of six germline AGOs that associate with 22G-RNAs: HRDE-1, PRG-1, WAGO-1, WAGO-3/PPW-2, WAGO-4, and CSR-1. We also surveyed the interaction networks of two RdRPs that synthesize 22G-RNAs, EGO-1 and RRF-1, and one germ granule helicase found in Z granules, ZNFX-1.Our results show that these proximal interaction networks reinforce the notion that PRG-1, WAGO-1, and WAGO-3/PPW-2 act in a shared regulatory pathway involving piRNAs and 22G-RNAs. In contrast, and despite sharing a set of 22G-RNAs that target an overlapping set of germline protein coding genes, CSR-1 and WAGO-4 have almost entirely distinct proximal protein interaction networks. The nuclear AGO, HRDE-1, reveals a somewhat surprising proximal interaction network involving splicing, transcriptional elongation, and termination factors, rather than chromatin modifiers. By overlapping all TurboID data with published TurboID datasets for two other core germ granule factors (DEPS-1 and GLH-1), we also identified a set of eleven largely uncharacterized, intrinsically disorder region (IDR)-domain possessing proteins that interact with multiple baits and contribute to fertility, germ granule structure/integrity and heritable RNA interference (RNAi). One of these proteins, D2005.4, appears to be a heme-binding protein that interacts with all of the germ granule AGOs. Based on its structure, encompassing a central Tetratricopeptide Repeat domain, flanked by two intrinsically disordered regions (IDR), and its TurboID protein interaction network, we speculate that this protein may be a scaffold for assembling RNA regulatory modules in the germ granules and may link germ granules to mitochondria. Overall, we anticipate that these TurboID datasets will serve as a rich resource for the field, and shine a light on unexpected functions of AGOs and RNAi factors.

## RESULTS

### Generating AGO Indirect TurboID *C. elegans* Strains

We prioritized the set of AGOs that localize to germ granules in young adult hermaphrodites, PRG-1, WAGO-1, WAGO-3/PPW-2, WAGO-4, and CSR-1b for these studies, because they are sufficiently expressed and predominantly localized to germ granules throughout the life cycle of the worm (Fig 1A)^3^. We also selected HRDE-1 as a comparator, which has been shown to localize to germ granules in its sRNA-unloaded state, yet is generally highly enriched in the nucleus^26^. Our lab recently generated and characterized a comprehensive set of *C. elegans* AGOs tagged with GFP::3xFLAG, which we crossed to a strain expressing a TurboID::antiGFP nanobody under the control of the *mex-5* germline promoter^3,33–35^. We confirmed that all TurboID/AGO strains displayed no defects in fertility (Fig S3), and that AGO localization was unaltered (data not shown) before examining biotin enrichment via streptavidin pulldown/blot experiments (Fig S1). Upon observing biotinylation patterns in the TurboID/AGO strains that were consistent with AGO expression, we proceeded to proximity labeling/mass spectrometry experiments on young adult hermaphrodites. In these experiments, the biotin source was from food, *E. coli OP50*, and therefore biotin labeling could occur at any stage in the worm lifecycle when both GFP::3xFLAG AGO and the TurboID nanobody were expressed. We performed experiments in triplicate, using an expression matched, germline expressed “TurboID nanobody-only” strain as well as wild-type (N2) worms with no TurboID nanobody nor GFP::3xFLAG::AGO as controls. We first compared all experimental data sets to N2 using SAINT (Significance Analysis of INTeractome)^36–38^ analysis to remove endogenously biotinylated proteins, and then used data from the TurboID nanobody-only strain to subtract non-specific labeling that could occur due to differences in AGO abundance. With this approach, we established a set of high confidence interactors for each AGO (Fig 1B).

**Figure 1.**
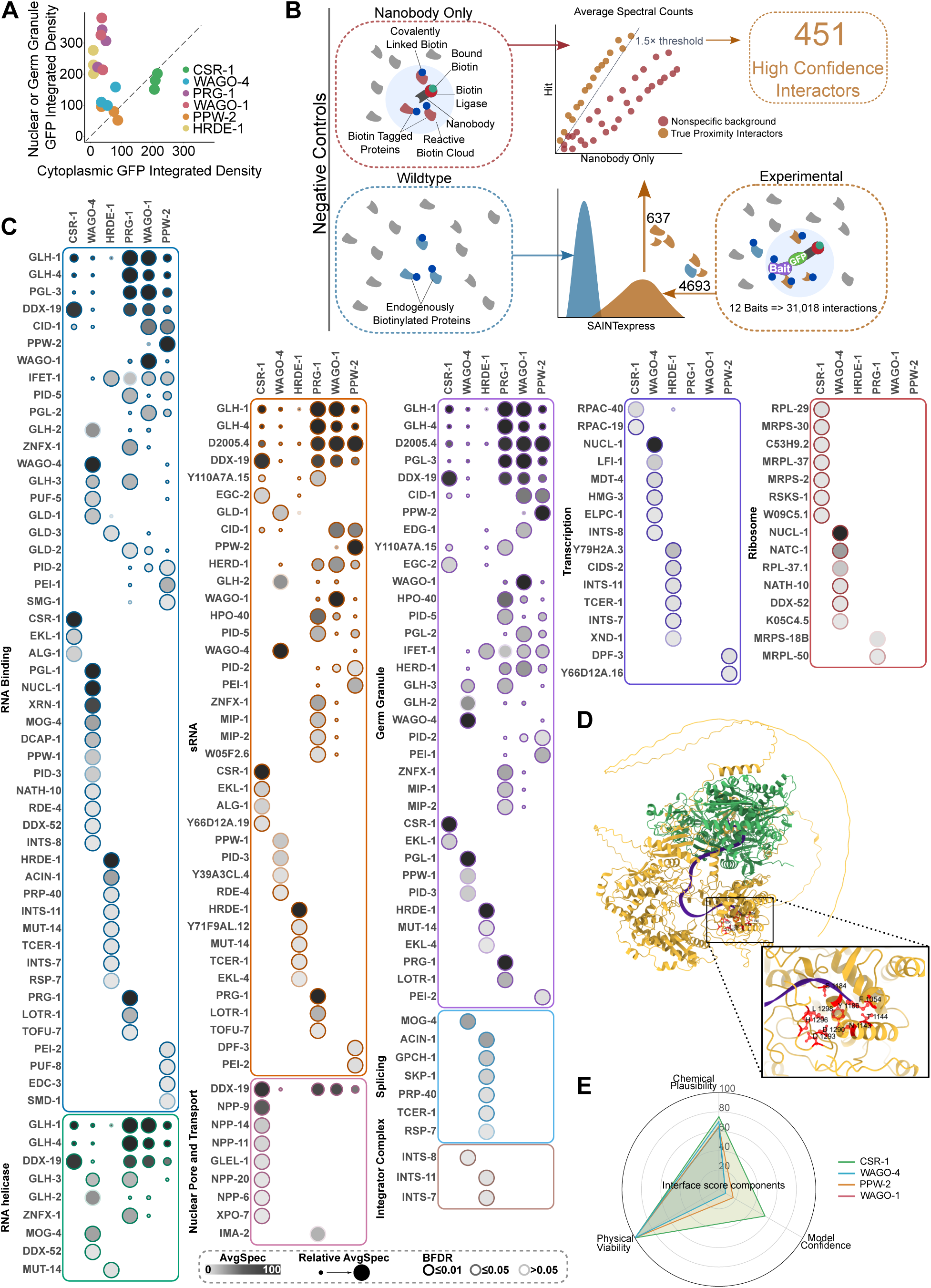
TurboID of germline AGO proteins reveals unique and overlapping networks. **A.** Fluorescence intensity of GFP-tagged AGO proteins was quantified in germ granules or nuclei compared to cytoplasm. Each datapoint represents a Region of Interest (ROI) including at least 20 nuclei within the pachytene region of a single adult hermaphrodite. **B.** In our approach, the TurboID biotin ligase is fused to a GFP nanobody that recognizes a GFP-tagged bait protein. TurboID enables the covalent linkage of biotin to nearby proteins, which are affinity-purified with streptavidin and subjected to mass spectrometry. Control strains include wild-type worms (N2), which have neither TurboID nor GFP-tagged AGO, and the TurboID expressing strain alone. SAINTexpress analysis was used to compare experimental samples to N2 controls, removing endogenously biotinylated proteins and generating interaction networks for each bait. Requiring 1.5 fold enrichment over the TurboID only control samples resulted in a high confidence interaction network for every AGO. **C.** Dot plot of all AGO TurboID data, in which the average spectral count is represented by node colour intensity, and the Bayesian False Discovery Rate (BFDR) by edge colour intensity. Proteins are grouped by manually curated sets of proteins, and because of overlap in these categories, a proximal interactor may be observed in more than one category. Additional protein categories are found in supplemental figures. **D.** Alphafold3 rendering of CSR-1 and CDE-1/CID-1 interaction, including an abundant 22G-RNA known to associate with CSR-1. Inset shows a zoom of the active site in CDE-1/CID-1 in association with the 22G-RNA 5′ end. **E.** Radar plot demonstrating the confidence in modelling the CDE-1/CID-1 interaction with CSR-1 and other AGOs that showed a proximal interaction.

### TurboID on Germline AGOs reveals unique and overlapping features of proximal protein interaction networks

The proximal interaction networks of each AGO highlight known and novel biological features unique to each AGO, and demonstrate the relationships between the AGOs. In this section, we highlight notable features of the proximal protein interaction networks of each AGO. In subsequent sections we examine the overlaps between proximal protein interaction networks of the AGOs with additional germ granule factors to gain insight into germ granule functions and organization.

### CSR-1b

CSR-1 consists of two isoforms, the long isoform, CSR-1a, which is expressed in spermatogenesis (L4) and the intestine, and CSR-1b, which is constitutively expressed in the germline. Because of the developmental timepoint and tissue we assayed in TurboID (young adult, when oocytes are made, and germline), our experiments examine CSR-1b (simply referred to as CSR-1 throughout). CSR-1 was initially characterized for its role in chromosome segregation in embryos in association with 22G-RNA biogenesis factors, DRH-3 (DEAD-box helicase), EGO-1 (RdRP), EKL-1 (Tudor domain protein), and CDE-1/CID-1 (terminal uridylyl-transferase)^19,39^. Consistent with this, we observed CSR-1 in association with EKL-1 and CDE-1/CID-1 in our TurboID datasets (Fig 1C and S4). To better understand if the interaction with CSR-1 and CDE-1/CID-1 was likely to be direct or indirect, we employed *in silico* modelling of these two proteins and a sRNA with AlphaFold3 (Fig 1D). In these studies, we observed that a region within an IDR of CDE-1 protrudes into the 3′ binding pocket of CSR-1, dislodging the 3′ end of the sRNA, which is displaced into the active site of CDE-/CID-1 (predicted according to homology to *S. pombe*^40,41^). This model seems plausible on the basis of CDE-1/CID-1’s role in uridylating the 3′ ends of CSR-1 22G-RNAs^39^. Notably, although multiple AGOs recover CDE-1/CID-1 as a proximal interactor (Fig 1C), the modeled interaction with CSR-1 displays the highest confidence and plausibility.

Among the AGOs analyzed, CSR-1 is the only AGO with a partial localization to D granules^13^. D granules are situated between the other germ granules and the cytoplasmic face of the nuclear pore, serving to anchor other germ granules to the nuclear periphery and enforcing the proper organization of granules^12^. Reduction in either CSR-1 or the cytoplasmic facing nuclear pore components NPP-10, -9, -20, and -7 results in dissociation of germ granules from the nuclear periphery^19,42^; loss of NPP-14 results in a reduction of CSR-1 and PGL-1^43^ recruitment to granules and loss of DDX-19 (a DEAD box helicase), or the nucleoporins GLEL-1, or NPP-14 results in aberrant germ granule mixing^12^. The association between nuclear pore proteins and D granule components is highlighted in the CSR-1 proximal interaction network, in which the nuclear pore proteins NPP-6, -9, -11, -14, and -20 and associated factors DDX-19 and GLEL-1 were identified as unique (not found in association with any other AGOs), high confidence proximal interactors of CSR-1 (Fig 1C). Of these nuclear pore proteins, NPP-9 and -14 are cytoplasmic filament proteins^43^, while NPP-6 and -20 are nuclear/cytoplasmic ring proteins^44^, all of which may contact germ granules. NPP-11 is a central channel protein^45^, and we speculate that this interaction may be indicative of CSR-1 trafficking through the nuclear pore, as CSR-1 has been reported in association with chromatin and nucleoplasm^46,47^. Consistent with this, Exportin 7 (XPO-7) was also identified as a CSR-1 proximal interactor. The additional identification of RNA Pol I/III regulatory subunits, RPAC-19 and RPAC-40, as proximal interactors also supports a nuclear presence for CSR-1 in the adult germline, although a role for CSR-1 in RNA Pol I/III transcription has not been reported so far. In addition to a proximal association with nuclear pore proteins, we observed an enrichment for the E granule component EGC-2, as well as conventional P granule components, GLH-1 and -4 (Vasa homologs), and PGL-3 (a ribonuclease)^5,31^ (Fig 1C). Our data reiterate a central role for CSR-1 acting in concert with nuclear pore proteins and D granule factors to maintain germ granule tethering and integrity, and highlight potentially novel nuclear roles for CSR-1.

A surprising finding from the CSR-1 proximal interaction data was the association of CSR-1 with proteins involved in metabolism and vesicular transport that are generally localized to the ER, Golgi apparatus, and mitochondria (Fig 1C and S4). In particular, mitochondrial proteins MRPS-2, MPRS-18B, MRPS-30, MRPS-37 stood out, as they are components of the mitochondrial ribosome. While the relationship between CSR-1 and cytoplasmic ribosomes has been established–for instance, our own TurboID data identify RPL-29 is a proximal interactor of CSR-1, and CSR-1 has been shown to promote 22G-RNA synthesis in conjunction with translation–the connection of CSR-1 to mitochondrial ribosomes has not been observed^48,49^. Given that mitochondrial ribosomes are resident on the inner mitochondrial membrane and encoded by the nuclear genome, and the fact that CSR-1 has not been observed to localize to the mitochondria by microscopy, we think it is likely that CSR-1 interacts with these proteins prior to their mitochondrial import. Collectively, these data clarify relationships between CSR-1 and its sRNA biogenesis factors, highlight links to the nuclear pore that are unique to this AGO, and point to yet to be discovered activities for this multi-functional AGO.

### WAGO-4

Of the full set of AGOs, WAGO-4 has a distinct relationship with CSR-1, because it binds to 22G-RNAs that target an overlapping set of germline expressed protein coding genes. In fact, >80 % of WAGO-4 targets are CSR-1 targets, yet WAGO-4 is thought to silence these targets, while CSR-1 licenses them^3,19,23,24,50^. Therefore, a major question in the field is how CSR-1 and WAGO-4 can be loaded with the same sRNAs and target the same genes, yet elicit different responses. Comparing the proximal interactomes of CSR-1 and WAGO-4 provides some clues. First, CSR-1 and WAGO-4 share very few proximal interactors, indicating that their roles in the germline are likely very different, despite possessing similar localization patterns and sRNA targets (Fig 1C). Second, the WAGO-4 proximal interaction network is enriched for proteins involved in mRNA stability and translational regulation, including DCAP-1, XRN-1, GLD-1, IFET-1, and PUF-5^51–55^ (Fig 1C and S4). The presence of these proteins in WAGO-4’s proximal interaction network, but not CSR-1’s suggests that WAGO-4, which lacks the catalytic tetrad of residues involved in endonucleolytic RNA cleavage, may induce mRNA decay (via DCAP-1 and XRN-1) and/or inhibit translation (via GLD-1, IFET-1, and PUF-5) rather than direct target cleavage, as WAGO-4 is thought to be a negative regulator of gene expression (Fig 1C).

WAGO-4 TurboID uncovered an abundance of helicases (Fig 1C). Several of these were shared with other germ granule AGOs (e.g., GLH-1, GLH-2* GLH-4, and DDX-19), while others were unique to WAGO-4 (e.g., DDX-52 and MOG-4). The connection between AGOs and helicases has been established across many species^56–58^. These interactions are prominent in germ granules, where Vasa and other DEAD/H box helicases have been implicated in the regulation of germ granule fluidity, transcript recruitment both *in vivo* and *in vitro*^59,60^, and sRNA biogenesis. WAGO-4 has been functionally and specifically linked to VASA helicases, as mutation or deletion of the FG repeats in GLH-1,-2, -4 (3xFG GLH mutant) resulted in a near complete loss of WAGO-4 from germ granules, but did not alter the association of CSR-1 or PRG-1 with germ granules^25^ (Fig 2A). The shift of WAGO-4 out of the germ granules and into the cytoplasm was accompanied by the loss of a subset of WAGO-4-specific 22G-RNAs (not shared with CSR-1), an enhanced loading of WAGO-4/CSR-1 shared 22G-RNAs into WAGO-4 complexes, and a downregulation of these shared mRNA targets relative to wild-type conditions^25^. We expected that examining the proximal interaction network of WAGO-4 in this 3xFG GLH mutant background would shed light on these molecular phenotypes, and aid in distinguishing germ granule-enriched vs. cytoplasmic interactions, so we performed TurboID on WAGO-4 in the 3xFG GLH mutant.

**Figure 2.**
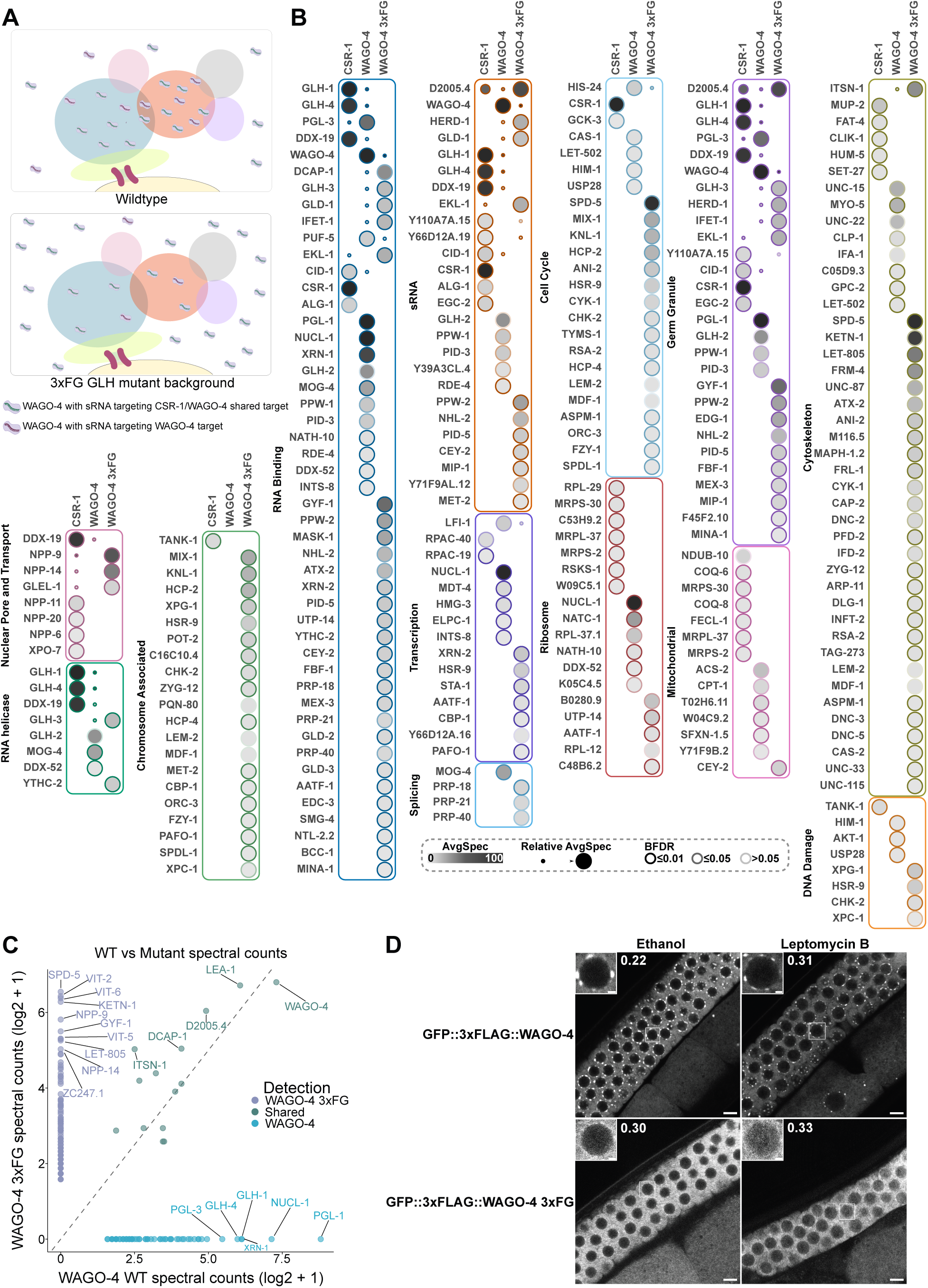
WAGO-4 displacement to the cytoplasm alters its interaction network. **A.** Diagram depicting wild-type WAGO-4 localization and its localization pattern in the 3xFG GLH mutant. Loss of FG repeats in three Vasa helicases (GLH-1, GLH-2, and GLH-4) leads to a decrease in WAGO-4 germ granule localization, and its displacement to the cytoplasm. This displacement results in alterations in bound 22G-RNA pools, with a decreased loading of WAGO-4 specific 22G-RNAs, and an increased loading of WAGO-4/CSR-1 shared 22G-RNAs, generated in the cytoplasm. **B.** Dot plot of CSR-1, WT WAGO-4, and 3xFG WAGO-4 TurboID data, in which the average spectral count is represented by node colour intensity, and the Bayesian False Discovery Rate (BFDR) by edge colour intensity. Proteins are grouped by manually curated sets of proteins, and because of overlap in these categories, a proximal interactor may be observed in more than one category. Additional protein categories are found in supplemental figures. **C.** Scatterplot comparing the log_2_(spectral counts +1) of interactors identified by mass spectrometry in WAGO-4 TurboID experiments in WT and 3xFG mutant backgrounds. Each point indicates a unique proximal interactor. Interactors with a log_2_(x+1) of 5 or greater are labeled. The dotted line indicates equal spectral counts between the two conditions. Note that baits were run on separate machines using different protocols TripleTOF6600 (WT) and timsTOF Pro 2 (3xFG), thus the scatterplot consists of spectral counts and not direct spectral comparisons. **D.** Worms expressing GFP::3xFLAG::WAGO-4 in WT and 3xFG mutant backgrounds were exposed to leptomycin B, or an ethanol vehicle control, at 20 °C for 4 hr before live imaging. Quantifications of nuclear:cytoplasmic ratios are shown beside nuclei insets. Scale bars represent 5μm and 1μm for nuclei insets.

We observed that the WAGO-4 interaction networks were very different between wild-type and the 3xFG GLH mutant, suggesting that we captured mostly germ granule-associated WAGO-4 in the wild-type experiments (Fig 2B,C, and S5). Consistent with WAGO-4’s enrichment in the cytoplasm, we saw that WAGO-4 lost interactions with all helicases except GLH-3, and gained an interaction with YTHC-2, an uncharacterized protein that has a DEAD-box helicase domain (Fig 2B). Remarkably, cytoplasmically-enriched WAGO-4 associated with an even greater number of proteins involved in post-transcriptional RNA regulation compared to wild-type. Proximal interactors involved in cap binding (DCAP-1, EDC-3)^61^, translation regulation (GYF-1, IFET-1)^54,62^ and 5′ end regulation coupled with RNA decay (XRN-2, SMG-4)^63,64^ were enriched, as were RNA binding proteins that typically associate with and regulate mRNA via the 3′ UTR, including PUF-5, FBF-1, GLD-1, GLD-3, NHL-2, Tag153/NTL-2.2 (Not-like) and MEX-3, as well as the polyA polymerase GLD-2^19,53,55,65–69^ (Fig 2B). This enhanced association with RNA decay factors when WAGO-4 is displaced from germ granules suggests that post-transcriptional regulation by this AGO and associated factors is likely to occur in the cytoplasm, and is consistent with the downregulation of shared CSR-1/WAGO-4 target transcripts in the 3xFG GLH mutant. In addition to these interactions, we observed an association between WAGO-4, MINA-1, and CEP-1/p53, substantiating the observation that MINA-1 and WAGO-4 genetically interact to control apoptosis and RNAi in the germline^70^.

Finally, WAGO-4 in the 3xFG background also associated with a large number of cell cycle, kinetochore, DNA damage repair, and cytokinesis proteins, which was surprising, given that WAGO-4 has not been observed in the nucleus before. We therefore more closely examined the localization patterns of WAGO-4 in the wild-type and 3xFG backgrounds, both under normal conditions and in the presence of leptomycin B, which blocks nuclear export. Under leptomycin B treatment, we identified a pool of WAGO-4 in the nucleus, with greater nuclear enrichment in the 3xFG background (Fig 2D). We speculate that these nuclear associations may underlie the endomitosis phenotypes observed in the 3xFG GLH mutant, in which oocytes aberrantly undergo repeated DNA replication in the absence of chromosome segregation elevated temperatures. Collectively, our TurboID data highlight a role for WAGO-4 in post-transcriptional regulation in the cytoplasm, reiterate the importance of helicases in recruiting WAGO-4 to the germ granules, and led us to uncover a heretofore unknown nuclear role for WAGO-4 in the germline.

### HRDE-1

HRDE-1 has been predominantly studied for its role in the nucleus, however, it was recently shown to cycle between the nucleus and SIMR foci^26^. In this cycle, in a non-sRNA bound state, HRDE-1 localizes to SIMR foci, where it is then loaded with sRNAs, inducing its migration to the nucleus to regulate its targets^71^. Many models suggest that, upon recognition of nascent RNA transcripts, HRDE-1 induces silencing by recruiting the H3K9 histone methyltransferases SET-25 and SET-32^72^. These methyltransferases deposit repressive histone modifications, which in turn recruit chromatin factors that lead to the formation of heterochromatin. More recent studies using tissue specific auxin-inducible degron systems support a model whereby HRDE-1 mediated silencing occurs independently of histone methylation^73^. Our HRDE-1 TurboID data support this model, as HRDE-1’s proximal protein interaction network did not reveal histone methyl transferases (Fig 1C and S4). Instead, our HRDE-1 interaction network was enriched for transcriptional and splicing factors that connect HRDE-1 to RNA Polymerase and implicate it in splicing regulation. First, and in common with CSR-1, we observed that RPAC-40, an Rpb3-like subunit of RNA Polymerases I and III, was a proximal interactor of HRDE-1. Like CSR-1, HRDE-1 has not been linked to regulation of RNAPol I or III transcripts, however, Alphafold structural predictions indicate the potential for RPAC-40 to interact with the Rpb-2 subunit of RNA Polymerase II (data not shown), suggesting that HRDE-1 could interact directly with any of the three polymerases via this factor. Several other proximal interactors link HRDE-1 to the largest subunit of RNA Polymerase II, which harbors the C-Terminal Domain (CTD), implicating HRDE-1 in the regulation of transcriptional termination and/or transcript 3′ end formation as well as splicing (see below). First, INTS-7 and INTS-11 are members of the Integrator complex, which processes the 3′ ends of snRNAs (U1, U2, etc.) and SL2 (Splice Leader RNAs), as well as some protein coding genes and piRNAs^74,75^ (Fig 1C). INTS-11 is an endonuclease that cleaves nascent RNAs, while INTS-7 is a structural subunit of the Integrator complex^75^. Second, CIDS-2, a largely uncharacterized protein with an RNA Polymerase II CTD interacting domain, has previously been linked to 3′ end cleavage, although it does not appear to possess any catalytic activity^76^. These data place HRDE-1 in direct or close contact with at least RNA Pol II, and possibly RNA Pol I and III.

Several proximal interactors link HRDE-1 to various steps in the splicing cycle and reinforce HRDE-1’s connection to the CTD (Fig 1C). Two proximal interactors, TCER-1 and SKP-1, link RNA Polymerase II and splicing^77–79^. TCER-1 associates with both the CTD and splicing factor SF1, which recognizes branch points during spliceosome assembly, bridging RNA Pol II and the spliceosome^80^. SKP-1 is homologous to human SKIP/SNW1, a component of the Prp19 complex that has links to transcriptional elongation via the P-TEFb complex and spliceosome activation later in the splicing cycle^81^. RSP-7 is an SR (Ser/Arg-rich) splicing factor involved in the snRNP recruitment, and whose loss results in P granule detachment from the nuclear periphery during embryogenesis, linking splicing to germ granule integrity^42,82^. PRP-40, is a component of the U1 snRNP, so far implicated in the inclusion of microexons in neurons, and active in the initial steps of splice site choice^83^. ACIN-1, a conserved component of the ASAP (apoptosis- and splicing-associated protein) complex, which is involved in alternative splicing and apoptosis, plays roles in the later steps of splicing and exon junction complex recruitment in humans^84^. These data are consistent with previous reports that HRDE-1 interacts with the conserved intron-binding helicase known as EMB-4/Aquarius and other nuclear RNA processing and export factors that act subsequent to splicing^85^.

While we did not observe histone methyl-transferases in the HRDE-1 TurboID dataset, we found several proteins with links to histone acetylation and genome stability (Fig 1C). Interactors with links to histone acetylation/chromatin remodeling include EKL-4 (homologous to human DMAP1, which could participate in NuA4 histone acetyl transferase/chromatin remodelling or SWR1 chromatin remodelling complexes)^86^ and the uncharacterized protein C16C10.4 (homologous to human SAP18, a member of the Sin3 histone de-acetylase complex)^87^. HRDE-1 proximal interactors with roles in genome stability touch on DNA replication and chromosome cohesion (CTF-4)^88^, repression of telomerase and alternative lengthening of telomeres (POT-2)^89^, and meiotic crossover distribution (XND-1)^90^.

Finally, we also captured several germ granule components (EDG-1, MUT-14, EGC-2) and RNA regulatory factors that may reflect cytoplasmic regulatory partners of HRDE-1, including the cytoplasmic polyA element binding protein, CPB-3, which induces 3′UTR shortening via alternative poly-adenylation (APA) in human cells^91^, along with other 3′ end RBPs including GLD-1 and GLD-3. In sum, the HRDE-1 TurboID data link this AGO more directly to the CTD of RNA Polymerase II rather than chromatin modulation via histone methyltransferases, and point to roles in transcriptional termination, 3′ end formation, and splicing.

### PRG-1 and WAGO-1

PRG-1, the only PIWI present in *C. elegans*, interacts with a class of genome-encoded sRNAs called piRNAs or 21U-RNAs to perform transcriptome surveillance^17^. piRNAs are transcribed by RNA Pol II as capped precursors from a cluster of piRNA loci on LGIV as well as from a set of protein coding genes, and then undergo a series of processing steps before being loaded into PRG-1 in the cytoplasm^20,92^. Here, piRNAs induce the production of 22G-RNAs that are loaded into WAGO class AGOS such as WAGO-1 and PPW-2/WAGO-3^3^. Consistent with this relationship, we observed a strong overlap in the proximal interaction networks of PRG-1 and WAGO-1 (60 % of hits shared, more below), and WAGO-1 was identified as a PRG-1 interactor (Fig 1C and S4). The PRG-1 and WAGO-1 interaction networks are enriched for helicases, RNA binding proteins required for Z and P granule organization and integrity, and several piRNA biogenesis pathway components, including PID-2 and PID-5^93^. PID-2 is an intrinsically disordered protein that works with the tudor domain/peptidase domain protein PID-5 to induce 22G-RNA silencing, perhaps physically linking PRG-1 to 22G-RNA biogenesis and WAGO loading. TOFU-7 was also identified in the PRG-1 proximal interaction network. TOFU-7 is a KH domain protein that is not well characterized, however, its loss leads to a depletion of piRNAs, and was speculated to act as a putative loading factor for PRG-1^94^. Additional components in the PRG-1 interaction network include uncharacterized proteins that may participate in ubiquitination, and proteins that are involved in post-transcriptional mRNA regulation, including IFET-1, GLD-2, and SMG-1^95^. Several other labs have also performed TurboID and our data are mostly consistent with their results. One unexpected aspect of the WAGO-1 proximal interactome was the enrichment of components of the DNA helicase MCM4/6/7 subcomplex, MCM-4 and MCM-6 (Fig 1C). Previous studies have shown that RNAi knockdown of MCMs causes dissociation of embryonic P granules^95^, and other groups using TurboID on germ granule factors have noted an enrichment for chromatin associated proteins, including kinetochore proteins like HCP-1, in phase separated cytoplasmic condensates^96^, suggesting the MCMs could behave similarly. Overall, our PRG-1 and WAGO-1 proximal interaction networks enrich for expected, as well as new interactors, and highlight the link between piRNA biogenesis and downstream 22G-RNA production.

### PPW-2

While PPW-2 is also thought to act in parallel to WAGO-1 downstream of piRNAs, and shares a large set of 22G-RNA targets with WAGO-1, the proximal interaction network of PPW-2 shares substantially fewer proteins with PRG-1 or WAGO-1 (Fig 1C and S4). PPW-2 has several well characterized interactors involved in paternal epigenetic inheritance, PEI-1 and PEI-2^21^, which were captured in the interaction network. These proteins are expressed during spermatogenesis and are present in spermatids contained within the adult hermaphrodite spermatheca. They are required for the formation of spermatogenic specific PEI granules and the retention of PPW-2 in spermatids^21^. Another well known interactor of PPW-2 is the dipeptidyl peptidase, DPF-3^97^, which also targets WAGO-1, yet DPF-3 was only identified as an interactor with PPW-2 here. Notably, PPW-2 identified WAGO-1 and WAGO-4 as interactors, along with a similar set of helicases (GLH-1, -3, -4), RNA binding (PGL-2, -3, PUF-5, -8) and post-transcriptional regulatory factors (EDC-3, IFET-1, SMG-1). In particular, the association of IFET-1, the eIF4E/IFE-3 transporter with multiple AGOs may indicate that these AGOs recruit or stabilize IFET-1 on target transcripts to block ribosome assembly and hold transcripts in stasis within germ granules. Like WAGO-1, PPW-2 enriches for PID-2 and PID-5, which are thought to connect piRNA silencing to 22G-RNA biogenesis^93^. Surprisingly, the PPW-2 proximal proteome is specifically enriched for vitellogenin/yolk proteins, VIT-1, VIT-3, and VIT-5. The proteins are made in the intestine and imported into the germline in order to provide nutrients to developing embryos. Recent evidence has found that yolk granules possess miRNAs capable of influencing larval gene expression^98^, though no evidence has found a connection to these proteins and 22G-RNAs, as such their presence in PPW-2’s proteome remains unclear. Overall, PPW-2’s interaction network points to both unique roles and overlapping functions with the other WAGOs expressed in the germline.

### Integration of AGO TurboID data into Germ Granule Proximal Interactomes reveals uncharacterized proteins as putative germ granule factors

To expand our understanding of how AGOs relate to germ granules, we overlapped AGO proximal proteomes with those of other germ granule factors from our lab (EGO-1, RRF-1, ZNFX-1) and several similar datasets of core granule proteins present in the field (GLH-1, DEPS-1)^30^ (Fig 3A and S6). Doing so enabled us to define a set of proteins that we term here the “core granule proteome:” proteins significantly enriched in three or more of the analyzed proteomes. In this section, we will discuss the individual proximal proteomes of ZNFX-1, EGO-1, and RRF-1, and then detail the largely uncharacterized proteins uncovered from overlapping these TurboID datasets.

**Figure 3.**
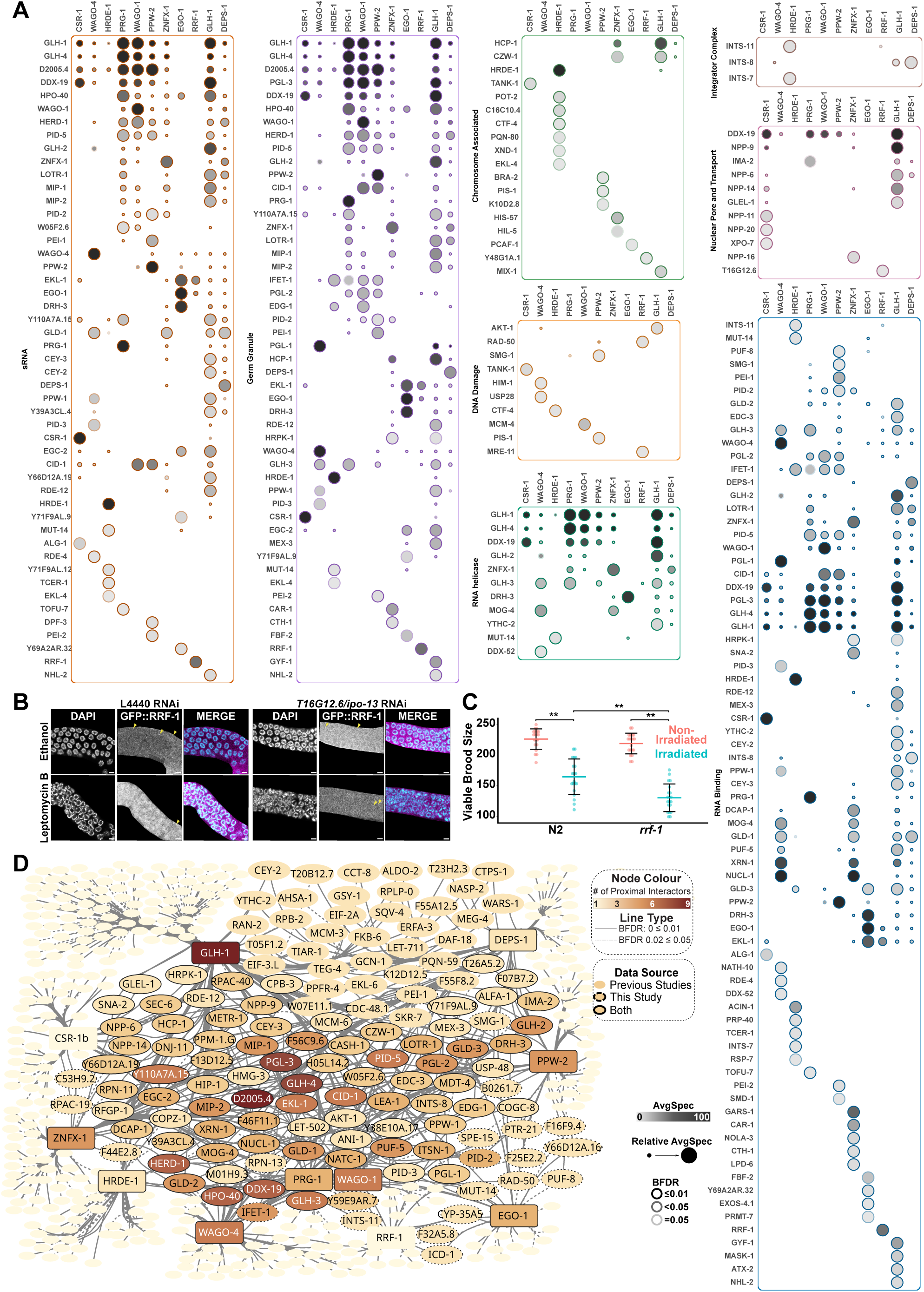
Expanding TurboID networks to other germ granule proteins reveals uncharacterized proteins. **A.** Dot plots of all WT AGO, ZNFX-1, RRF-1, EGO-1, DEPS-1, and GLH-1 TurboID data, in which the average spectral count is represented by node colour intensity, and the Bayesian False Discovery Rate (BFDR) by edge colour intensity. Proteins are grouped by manually curated sets of proteins, and because of overlap in these categories, a proximal interactor may be observed in more than one category. Additional protein categories are found in supplemental figures. **B.** Worms expressing GFP::RRF-1 were exposed to either empty RNAi vector (L4440) or *ipo-13* (*T16G12.6*) RNAi, then subjected to treatment with leptomycin B, or an ethanol vehicle control, at 20 °C for 4 hr before fixation, staining with DAPI, and imaging. Yellow arrowheads point to GFP::RRF-1 in Mutator Foci. Scale bar represents 20μm. **C.** WT (N2) or *rrf-1* young adult worms were exposed to 60 Gy γ-rays using a Caesium-137 source, then viable progeny were counted. N = 20 worms per condition. Significance was determined with one-way ANOVA multiple comparison test followed by post-hoc analysis by Tukey’s test. ** indicates significance of *P*< 0.01 **D.** Cytoscape map of all WT AGO, ZNFX-1, RRF-1, EGO-1, DEPS-1, and GLH-1 TurboID data, Only preys identified in at least two datasets are labeled. The number of identifying baits for each prey is indicated via colour shift from light yellow to dark red and the Bayesian False Discovery Rate (BFDR) by solid (less than or equal to 0.01) or dashed (between 0.02 and 0.05) edges.

### RRF-1 and EGO-1

EGO-1 and RRF-1 are partially redundant RdRPs responsible for 22G-RNA synthesis^19,20,99^. Although both RdRPs display the lowest-expression of all baits used in this study, both identified themselves as their most enriched prey, indicating sufficient targeting of the TurboID nanobody and accurate labeling. EGO-1 and RRF-1 play distinct roles in sRNA pathways, which is reflected in the proximal proteomes of each RdRP. EGO-1 is responsible for the majority of 22G-RNA synthesis, including both the CSR-1 and WAGO pathways, and is reported to reside in E granules^11^, while RRF-1 contributes a subset of WAGO class 22G-RNAs and resides in Mutator foci^8^. The EGO-1 proximal proteome is enriched for RNA binding proteins (PUF-8, MEX-3, FBF-2, and GLD-3) as well as post-transcriptional regulators (GLD-2, EXOS-4.1) (Fig 3A). The EGO-1 proteome is also enriched for the AGOs PPW-2 and WAGO-4, which could indicate that AGOs associate with the RdRPs to facilitate sRNA loading, a poorly understood step in the 22G-RNA pathway. EGO-1 also showed a proximal interaction with the DEAD box helicase MUT-14, found in Mutator foci. Both EGO-1 and RRF-1 were previously found to genetically and physically interact with the dual tudor domain protein, EKL-1 and the DEAD box helicase DRH-3^20^. EKL-1 was enriched in both the EGO-1 and RRF-1 proximal proteomes, while DRH-3 was only identified in the RRF-1 proteome. The RRF-1 proximal interaction network is also enriched for WAGO-4, but not PPW-2, along with the RNA binding proteins GLH-3, PGL-2, and the decapping scaffold, EDC-3 (Fig 3A). Surprisingly, the RRF-1 proteome enriched for EGO-1, but not vice versa, indicating that there may be cooperation or competition between the RdRP complexes and co-factors. Another surprising finding was the interaction of RRF-1 with members of the double strand DNA break (DSB) repair pathway MRN complex components MRE-11 and RAD-50^100^, as well as the endonuclease of the Integrator complex, INTS-11, which was also enriched in the HRDE-1 interaction network, and a nuclear transporter/karyopherin IPO-13 (T16G12.6)^101^. These findings suggest additional nuclear roles for RRF-1 that have not been described. To test for nuclear localization of RRF-1, we treated GFP::RRF-1 expressing worms with leptomycin B, and observed RRF-1 in the nucleus (Fig 3B). Next, we repeated this experiment, with the addition of RNAi against *ipo-13*. In this experiment, we found that RRF-1 was not as robustly enriched in the nucleus under *ipo-13* RNAi (Fig 3B), suggesting that IPO-13 is responsible for at least some of the nuclear import of RRF-1. To test for a role for RRF-1 in the DSB pathway, we subjected WT and *rrf-1* mutant worms to gamma irradiation, which induces DSBs and examined viable brood size. Mutants of *rrf-1* displayed a significant reduction in viable brood size relative to irradiated WT and non-irradiated controls (Fig 3C). Taken together, these data point to important and unappreciated nuclear roles for RRF-1. Overall the proximal interaction network of EGO-1 yielded known and expected interactors, while the interaction network of RRF-1 provided key insights into potential nuclear role(s).

### ZNFX-1

The helicase ZNFX-1 was first characterized in association with WAGO-4, as the two defining members of Z granules^10^. ZNFX-1 is involved in transmitting sRNAs during TEI, and its loss leads to a decrease in 22G-RNAs at the 3′ end of transcripts, along with decreased pUGylation^102^. We performed both direct and indirect TurboID on ZNFX-1 and combined the results to generate a robust and comprehensive proximal interaction network. The proximal interaction network for ZNFX-1 is indicative of its keystone position within the germ granule hierarchy, as a protein involved in coordinating multiple sRNA pathway activities, with a clear enrichment of RNA binding proteins, helicases, and sRNA pathway proteins (Fig 3A and S6). From the sRNA pathway standpoint, we observed EKL-1, DRH-3, and CID-1/CDE-1, all components of the CSR-1 22G-RNA biogenesis pathway. PID-2 and PID-5 were also present, along with PRG-1 and WAGO-1, connecting piRNA biogenesis to WAGO 22G-RNA synthesis and TEI. Although ZNFX-1, EGO-1, and RRF-1 have been shown to localize to distinct perinuclear foci, Z granules, E granules, and Mutator foci, respectively, all three proteomes identify known components across the various germ granule subcompartments. The discovery of proteins across germ granules in these three proteomes emphasize the potential and likelihood that many proteins move from one subcompartment to another.

### Overlapping proteomes reveal uncharacterized proteins with potential to act in germ granule and sRNA biology

We next integrated all of our TurboID datasets (ZNFX-1, EGO-1, RRF-1, CSR-1, WAGO-4, HRDE-1, PRG-1, WAGO-1, PPW-2) with two previously generated datasets for key P granule components, GLH-1 (helicase) and DEPS-1 (germ granule nucleating protein/RNA binding protein), which we re-analyzed using SAINT to be able to compare the data directly^36–38^. In doing so, we uncovered a high confidence “core” set of known and likely germ granule proteins that overlap in three or more of the eleven TurboID interaction networks (Fig 3D and S6). Germ granules across species are characterized by three major types of proteins: helicases (e.g. GLH-1, -2, -4, ZNFX-1, etc.), tudor domain proteins (EKL-4, LOTR-1), and AGOs, which are all abundantly represented across datasets. Canonical P granule markers PGL-1 and PGL-2 are broadly enriched, as are biogenesis factors for multiple sRNA pathways, including RdRP complex members (e.g., EKL-1 and DRH-3), AGOs (e.g., WAGO-1 and PRG-1), sRNA modification machinery (e.g., CID-1, TOFU-2), as well as a number of different RNA binding proteins, mRNA modification machinery, and mRNA degradation machinery. As anticipated, ribosomal proteins are largely depleted, except from CSR-1 and WAGO-4 data sets, indicating interactions in the bulk cytoplasm. In addition to these previously identified germ granule and sRNA pathway proteins, our analysis highlighted a high confidence set of potential germ granule proteins that had not been studied in depth previously. These factors include D2005.4, Y39A3CL.4, C27B7.5 (EGC-2), Y111B2A.3 (HERD-1), Y110A7A.15, F13D12.5 (ROOM-2), F46F11.1, F56C9.6 (EPS-1), Y66D12A.19, M01H9.3, and W05F2.6 (Fig 4A). These novel hits, while still largely uncharacterized, are not entirely new to the field, and many are present in other proteomics datasets. Several have also been identified via mass spectrometry and proximity labeling experiments for other germ granule factors, and throughout the duration of this project, some were characterized to varying extents, indicating that they are robust interactors of germ granules^31,43,103,104^.

**Figure 4.**
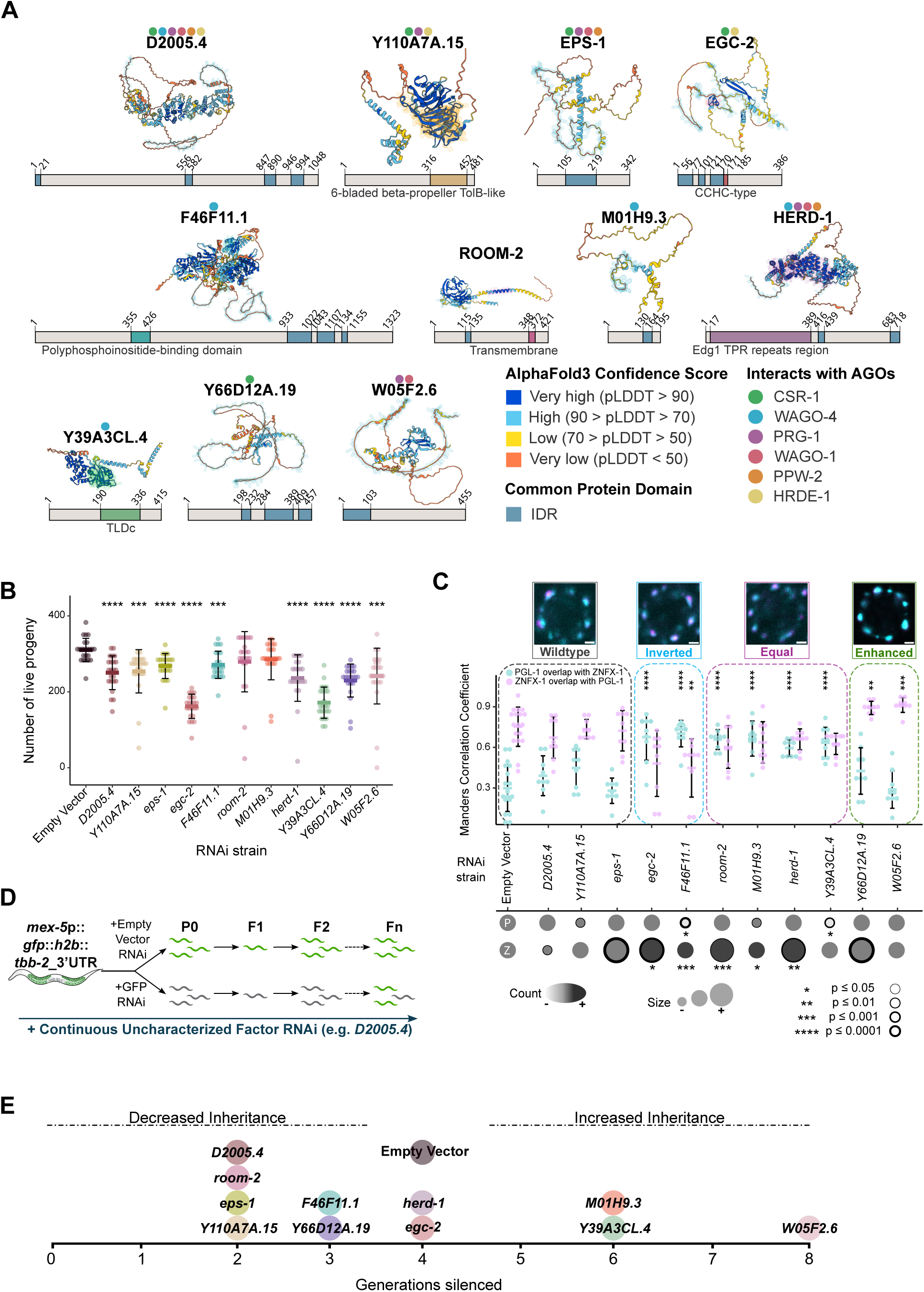
Uncharacterized proteins play roles in germ granule physiology, fertility, and RNAi. **A.** AlphaFold database predicted structures and linear domain maps for 11 proteins of interest. Each structure is coloured according to the Predicted Local Distance Difference Test (pLDDT), with blue being highest confidence, and orange lowest confidence. Domains from the corresponding linear protein map are also highlighted on the structure prediction. Intrinsically disordered regions (IDRs) are a common domain across the majority of the proteins and shown in blue. Coloured dots above the protein name correspond to AGO networks in which the protein was a prey. **B.** Brood size of worms fed RNAi against the genes of interest as well as empty vector (L4440) as a control. Significance was calculated via t-test. ***p ≤ 0.001, **** p ≤ 0.0001, N => 20 worms for each treatment. **C.** Germ granules of live RNAi-fed PGL-1::BFP; ZNFX-1::RFP worms were analyzed for colocalization, size, and number. PGL-1 is a marker representing P granules, and ZNFX-1 represents Z granules. Channels were aligned using TetraSpeck microspheres in FIJI and manually thresholded to distinguish germ granules (see Methods). Manders Correlation Coefficient was measured for PGL-1 signal overlap in ZNFX-1 and vice versa. Granule count and size were measured using the Analyze Particles feature in FIJI. Significance was calculated using a t-test against empty vector of the corresponding measurement. **D.** Outline of the Transgenerational Epigenetic Inheritance (TEI) assay. Worms treated with RNAi against the genes of interest for the entirety of the assay were also treated with anti-GFP or control RNAi for a single generation (P0). Successfully GFP-silenced worms were single-picked to their corresponding RNAi and the line was monitored for the re-expression of GFP in the germline of at least 75 % of the worms. **E.** Number of generations silenced in the TEI assay. RNAi treatments resulting in less than 4 silenced generations were categorized to have decreased inheritance whereas those that were silenced beyond 4 generations were categorized to have increased inheritance as per the performance of the empty vector (L4440) control. Worms were considered to be “ON” in generations that showed at least 80 % of the worms regaining GFP expression.

We first examined the predicted structures of these proteins using AlphaFold3, and noted that nearly all displayed intrinsically disordered and low complexity regions, which are common among proteins involved in phase separation (Fig 4A). In addition to IDRs several proteins contain domains that may hint at their function. D2005.4 possesses an internal Tetratricopeptide Repeat region (TPR), commonly found among chaperones and scaffold proteins to mediate protein-protein interactions^105^. HERD-1 also possesses a well-defined TPR region. F46F11.1 has homology to human VIP1 and possesses a Polyphosphoinositide-binding domain, which may mean that it is recruited to membranes formed between nuclei in the syncytial germline (called intercellular bridges). Similarly ROOM-2 contains a transmembrane domain and associates with intercellular bridges, suggesting that its identification in germ granule related datasets may involve contact between granules and these membranes. Finally, Y39A3CL.4 possesses a Tre2/Bub2/Cdc16 (TBC), lysin motif (LysM) domain catalytic (TLDc) domain. Though the exact molecular mechanisms of this domain remain poorly understood, much of what is known comes from mammalian cell systems where proteins with these domains are involved in protecting the cell from reactive oxygen species^106^. In addition to structural analysis, we examined the conservation of these proteins across nematodes and found that the majority of these factors are reasonably well-conserved across the *Caenorhabditis* genus (Fig S7). Some, such as F46F11.1 and Y39A4CL.4, are more broadly conserved across *Nematoda* as a whole. Interestingly, W05F2.6 stands out because it appears to be unique to *C. elegans.* With these putative germ granule factors in hand, we set out to understand how they may contribute to germ granule formation and function.

### Loss of putative germ granule factors leads to changes in fertility, RNAi inheritance, and germ granule morphology

For efficiency in examining a variety of phenotypes across this set of eleven proteins, we assayed the loss of function phenotypes of the uncharacterized factors using RNAi. Because germ granule factors play roles in fertility, RNAi, epigenetic inheritance, and the formation and maintenance of germ granules themselves, we assessed each of these features. First, we assayed viable brood size at 20 °C upon RNAi knockdown of each factor to determine if they play roles in fertility (Fig 4B). We observed that depletion of *C27B7.5 (EGC-2) or Y39A3CL.4* led to the most severe reduction in fertility (48 % and 45 % decreases in fertility relative to wild type/N2, respectively), loss of *Y66D12A.19, Y111B2A.3 (HERD-1),* or *W05F2.6* displayed moderate reductions in fertility (26 %, 24 %, and 22 % decreases, respectively), and loss of *D2005.4, Y110A7A.15, F56C9.6 (EPS-1)*, or *F4611.1* resulted in mild reductions in fertility (19 %, 18 %, 14 %, and 13 % decrease, respectively). *F13D12.5 (ROOM-2)* and *M01H9.3* showed no significant impact on fertility in this assay.

Next, because loss of germ granule and sRNA pathway components can impact germ granule formation, stability, and morphology, we examined whether loss of these uncharacterized factors leads to changes in size, number and overlap between the well characterized P and Z granule components, PGL-1 and ZNFX-1, respectively, using Mander’s correlation coefficient (Fig 4C). Normally, P granules are larger than Z granules, and the overlap in pixels between PGL-1 with ZNFX-1 is lower (i.e., less of the P granule overlaps with the Z granule) than the ZNFX-1 overlap with PGL-1 (i.e., more of the Z granule overlaps with the P granule). While a few of our uncharacterized factors showed no significant impact on granule size or morphology, most showed a range of phenotypes. Remarkably, loss of most factors, except for *D2005.4, Y110A7A.15, and Y39A3CL.4,* resulted in an increase in the number of Z granules. Conversely, loss of only *D2005.4* or *HERD-1* led to an increase in the number of P granules. Loss of *D2005.4* and *Y110A7A.15* led to no change in the size or morphology of the granules. Similarly, loss of *EPS-1* resulted in no impact on their relative pixel overlap between PGL-1 and ZNFX-1, yet led to larger P and Z granules, suggesting that it plays a role in regulating the size but not the separation of these germ granule components. Loss of *EGC-2* and *F46F11.1* led to an “inverted” granule phenotype, in which there was greater overlap of PGL-1 with ZNFX-1, accompanied by increases in Z granule number. Loss of *ROOM-2, M01H9.3, HERD-1,* and *Y39A3CL.4* resulted in an equalization in the overlap of P and Z granule pixels, suggesting that their loss led to de-mixing of the granules (“equal” phenotype), while loss of *Y66D12A.19,* and *W05F2.6* resulted in the partial or complete envelopment of P granules by Z granules and an increase in Z granule numbers. Collectively, these factors appear to play roles in germ granule size, number, and the extent of mixing between P and Z granule components. These phenotypes are generally consistent with fertility outcomes, wherein those with unaffected or “equal” germ granules displayed brood sizes closer to the control, while those with more extreme germ granule phenotypes (“inverted” or “enhanced”) had more extensive brood decreases.

Finally, we examined a role for these factors in epigenetic inheritance. To do this, we tested RNAi initiation and inheritance, using a germline GFP transgene and RNAi against both GFP or an empty vector and the uncharacterized factors^107^. We initiated the GFP/empty vector RNAi treatment in the P0 generation to test for initiation of an RNAi response. In the F1 and subsequent generations, we omitted the GFP/empty vector RNAi, and only knocked down the uncharacterized factors, to determine if they are involved in the inheritance of RNAi (Fig 4D). Normally, RNAi can perpetuate for four to five generations after the initial dsRNA trigger^108^. Deviations from this timing are indicative of factors that are involved in inheritance, either as “brakes” on inheritance (in which case, loss of the factor results in prolonged RNAi inheritance) or as factors that are required for the inheritance of RNAi (in which case, loss of the factor results in RNAi inheritance for fewer generations). Worms in which the uncharacterized factors were knocked down were still able to initiate GFP RNAi, indicating they do not play a role in the initial silencing response (i.e., they are not RNAi Deficient, Rde) (Fig 4E). Loss of two of the factors (*EGC-2* and *HERD-1)* appeared to have no impact on TEI. In contrast, the majority of uncharacterized factors did significantly alter epigenetic inheritance of RNAi silencing. Knockdown of six out of the eleven uncharacterized factors (*Y110A7A.15, EPS-1, ROOM-2, D2005.4, Y66D12A.19,* and *F46F11.1*) resulted in decreased inheritance with the most severe decreasing being *Y110A7A.15, F56C9.6, ROOM-2,* and *D2005.4* lasting only two generations. RNAi against the remaining three factors (*Y39A3CL.4, M01H9.3,* and *W05F2.6*) resulted in extended silencing by several generations.

The most pronounced effect was from loss of *W0F52.6,* which extended RNAi perdurance twice as long as the empty vector treatment (i.e., to 8 generations). Our results point to roles for these factors in TEI and RNAi, yet further experimentation will be necessary to determine their exact roles in these processes. Altogether, these experiments point to roles for the uncharacterized proteins in fertility, germ granule number, size and morphology and TEI/sRNA pathways, highlighting the utility of TurboID to identify new germ granule, fertility, and RNAi factors.

### D2005.4: A new germ granule factor with potential for scaffolding and enzymatic activity

Of all the uncharacterized germ granule factors we identified, D2005.4 stood out, as it was enriched in every AGO TurboID dataset (Fig 1C), as well as in association with ZNFX-1, DEPS-1, and GLH-1 (Fig 3A). D2005.4 encodes an uncharacterized protein with N and C terminal IDRs flanking a putative TPR central domain. Recent reports showed that D2005.4 localizes to germ granules and indicates a role for D2005.4 in sRNA biogenesis, as its loss leads to a depletion of 3′ piRNA dependent and independent 22G-RNAs^32^. In our own characterization of D2005.4 by RNAi knockdown, we observed a decreased brood, a decrease in the size of Z granules, and reduced RNAi inheritance (Fig 4B, C and E). To better characterize the role of D2005.4 in sRNA related pathways and define D2005.4’s protein interaction network, we again turned to TurboID, using an existing C-terminally GFP tagged D2005.4 strain^13^.

The proximal interaction network of D2005.4 reflects the potential for different roles for D2005.4 in different subcellular compartments. There was an enrichment in RNA binding and regulatory proteins (FBF-1, GLD-2, PGL-2, CEY-2, GLH-3), and sRNA biogenesis factors (TOFU-2, MUT-7, CID-1), reflecting roles in germ granules and sRNA pathways (Fig 5A). In addition, our data indicate a potential role for D2005.4 in association with the mitochondria, as another major category of interactors were mitochondrial-localized enzymes and proteins. Interestingly, D2005.4 recovered D2005.7, a small protein predicted to be associated with mitochondria encoded by the gene just downstream of D2005.4, as a proximal interactor. Other noteworthy categories of D2005.4 proximal interactors were cytoskeletal proteins, secretory/vesicular proteins, and stress response proteins. Somewhat surprisingly, the proximal interaction network of D2005.4 did not show enrichment for AGOs.

**Figure 5.**
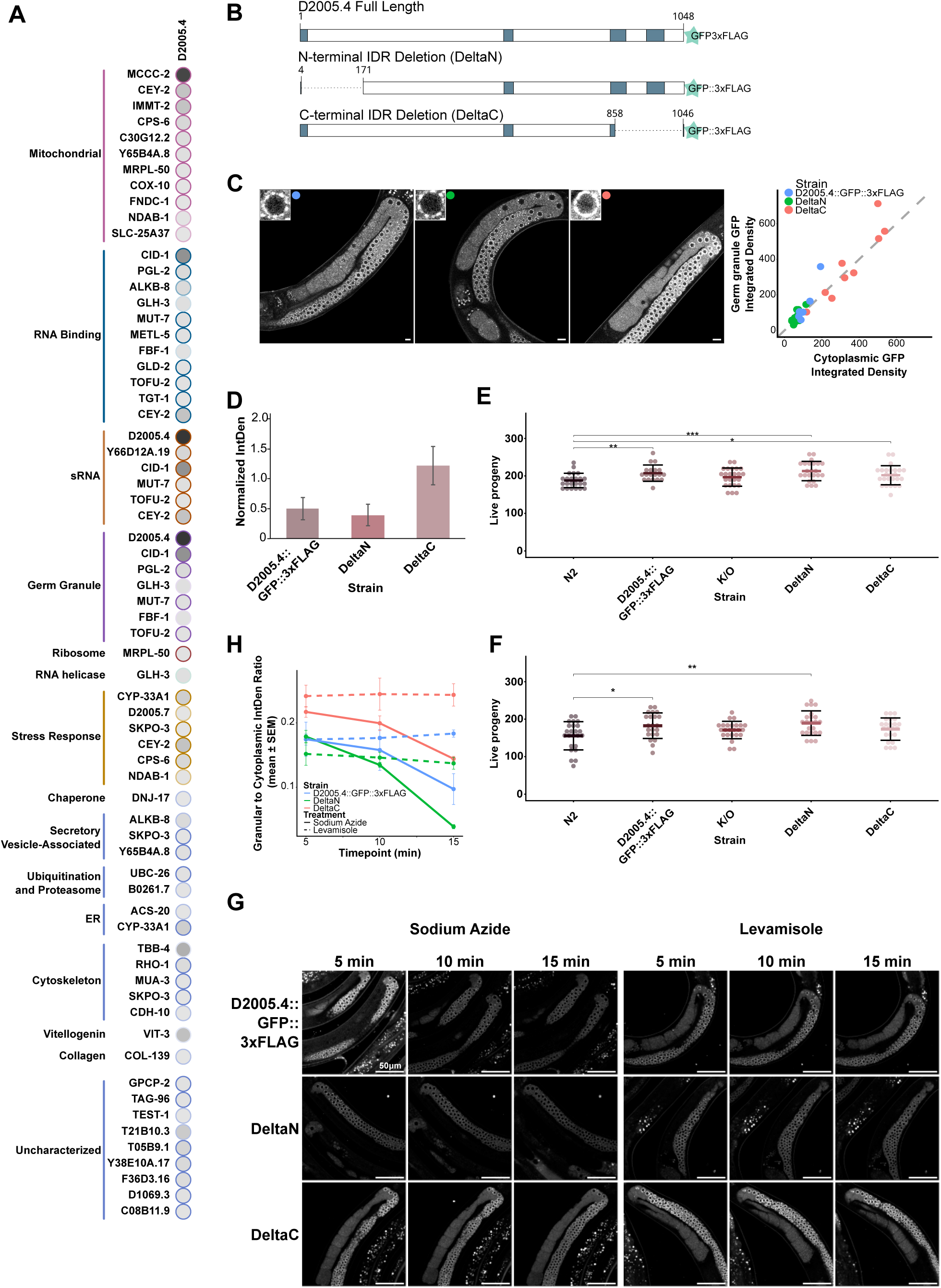
D2005.4 is a heme binding germ granule protein. **A.** Dot plot D2005.4 TurboID data, in which the average spectral count is represented by node colour intensity, and the Bayesian False Discovery Rate (BFDR) by solid (less than or equal to 0.01) or dashed (between 0.02 and 0.05) edges. Proteins are grouped by manually curated sets as marked. **B.** Domain diagram of *D2005.4* and mutants generated. Intrinsically disordered regions (IDRs) in blue, based on UnitProt predictions. Deletion mutants were designed using predicted Alphafold3 structures to guide IDR boundaries. **C.** Live imaging of animals expressing D2005.4::GFP::3xFLAG (WT), D2005.4::GFP::3xFLAG (DeltaN), and D2005.4::GFP::3xFLAG (DeltaC). Scale bar represents 10 µm for full germline and 1 µm for single nuclei insets. Granule vs. cytoplasmic signal was determined by relative thresholding and integrated densities plotted against each other. Each point represents a separate animal. **D.** Quantification of dilution series for D2005.4::GFP::3xFLAG (WT) and the domain mutants. Dilutions consisted of lysate equating to 100, 50, and 25 worms for each strain (N=2). Original western blot can be found in Supplemental Figure S8. **E.** Brood size assay of all D2005.4 strains compared to N2 conducted at 20 °C and 25 °C. 25 animals/condition were initially, only counts from animals that survived the assay were included. Each point consists of the entire brood size of a single animal. **F.** Mortal germline assay for GFP tagged Wildtype D2005.4 (WT) and the domain mutants compared to N2. Synchronized worm populations were plated at 25 °C and four L4 animals were transferred every three days. A total of 10 plates per condition were used to start the assay, a plate was considered dropped when no progeny was present to be transferred. **G.** Time lapse imaging of D2005.4 (WT) compared to domain mutants on sodium azide and levamisole. All imaged worms are gravid adults from synchronized populations. Azide treated worms received 20mM sodium azide. Levamisole treated worms received 2mM Levamisole. Quantification consists of replicates (n=3) imaged on separate slides, time count begins from the moment worms are picked into the immobilization agent. **H.** Time course quantification of D2005.4::GFP fluorescence in WT, DeltaN, and DeltaC backgrounds treated with either 20 mM NaN_3_ (solid lines) or 2mM levamisole (dotted lines).

IDRs have been implicated in recruiting proteins to phase separated condensates. To test the role of D2005.4’s N and C terminal IDRs in its recruitment to germ granules, and other functions, we made separate deletions of the N terminal IDR and the C terminal IDR (referred to as DeltaN and DeltaC), along with a null mutant that eliminates the full open reading frame (Fig 5B). We first examined protein levels and localization of these mutant versions of D2005.4. Loss of either IDR does not dramatically change the localization of D2005.4 to germ granules, however levels of the N terminal IDR deletion protein are slightly lower, while the C terminal deletion protein is slightly higher than WT (Fig 5C and D, Fig S8). These observations indicate that the IDRs are not the main drivers of D2005.4’s association with germ granules. Surprisingly, deletion of the full *D2005.4* coding region did not result in a substantial decrease in brood size at 20 °C, nor at 25 °C (as was previously reported^32^) (Fig 5E and F). However, this strain displayed a mortal germline phenotype after 10 generations at 25 °C (Fig S8), indicating a transgenerational epigenetic role (Fig S8). Further, when we performed RNAi against both *D2005.3*, the gene immediately upstream of *D2005.4*, in the *D2005.4* deletion strain at 25 °C, we observed a reduced brood size (Fig S8). Because of the close proximity of the 5′ ends of both genes, it may be that deleting the first intron or even the 5′ coding regions of *D2005.4* could alter *D2005.3* expression, leading to a loss of function for both genes and resulting in a reduced brood size at 25 °C.

Somewhat serendipitously, we identified a role for heme in D2005.4 stability. While performing live imaging using sodium azide to paralyze the worms we noticed that the D2005.4::GFP signal was reduced over the course of the imaging session (Fig 5G and H). Sodium azide chelates iron from heme, rendering heme binding proteins, including those of the mitochondrial electron transport chain non-functional. Importantly, we did not observe destabilization of D2005.4 when using the paralytic agent levamisole which causes muscle depolarization, or with CCCP (Carbonyl cyanide m-chlorophenyl hydrazone, data not shown), which uncouples the electron transport chain (thus ruling out a mitochondrial effect). We used AlphaFold3 to model the structure of D2005.4 with and without heme, and found that the addition of heme improved the confidence of the structural prediction overall (data not shown). While the molecular role of D2005.4 in germ granules, sRNA pathways, and mitochondria remains to be fully determined, based on the Turbo ID networks it was identified in and its own set of proximal interactors, it is seems likely that this protein may play important scaffolding and possibly enzymatic roles in these pathways.

## DISCUSSION

IP-MS has been a powerful tool to identify protein-protein interactions, however it has several significant challenges, including the potential for artifacts to arise in the preparation of lysates when proteins that normally reside in separate cellular compartments are mixed and reduced potential to recover transient or weak interactions. In contrast, *in vivo* labelling approaches such as TurboID have the capacity to uncover proteins in physical proximity within living cells, including dynamic or low-affinity interactions that may still be biologically meaningful. Here, we applied TurboID to define the proximal proteome of germline AGOs enriched in germ granules, along with the RdRPs RRF-1 and EGO-1, and the helicase ZNFX-1. Comparing these datasets, and to TurboID data for other germ granule proteins GLH-1 and DEPS-1, enabled us to uncover new insights about the functions of sRNA pathways and allowed us to identify a set of uncharacterized proteins that appear to play roles in germ granule biology, including the likely heme-binding scaffold, D2005.4. By viewing our TurboID approach as a proteomic “screen” and using various *in vivo* and Alphafold modelling approaches to verify unexpected interactions, we uncovered new localization patterns and activities for RNA pathway components (WAGO-4, HRDE-1, RRF-1), and explanations for how longstanding pathway partners interact (CSR-1 and CID-1/CDE-1).

### Considerations for using TurboID

Several features of our approach to TurboID bear noting, as they influence the conclusions we are able to make from these data. Because we did not deplete worms of biotin, nor did we temporally restrict when TurboID was expressed, aside from using a tissue specific germline promoter, labelling could occur in the germline at any point in the life cycle of the worm. This means that the proteins captured in our assays could have been interactors at different developmental stages, depending on the turnover of the labelled proteins. This also means that endogenously-biotinylated proteins are biotinylated as they normally would be, and the worms remain healthier overall. The use of SAINT analysis allows us to determine the likelihood an identified hit is a false-positive and thus allows for a more stringent determination of which proteins are true hits. Using N2 possessing no TurboID nor GFP-tagged protein, and nanobody-only negative controls for all experiments provided a rich and extensive source of comparative data that enabled the subtraction of endogenously biotinylated proteins with high confidence. Comparison of hits across the datasets also enhances confidence in the results, because we observed consistent hits that fit with germ granule localization (e.g. GLH proteins) across multiple baits, alongside unexpected hits that are specific to different AGOs (e.g., MCM proteins with WAGO-1). If these unexpected interactors were simply technical artifacts of abundant proteins or a result of the location of the GFP::TurboID nanobody, we would expect to observe them across datasets, including controls.

The position of the GFP tag in the bait protein is another consideration that influences the recovery of interactors and the interpretation of results. Because of the size of GFP and the nanobody, we calculate that the labeling radius for indirect TurboID is likely closer to 20 nm, rather than the 10 nm cited in the literature for direct TurboID. As with direct TurboID, placing the enzyme at the N or C terminus of the protein (or even in the middle, for that matter), could lead to the labeling of proteins that associate with one part of the protein or the other, yielding differing results.

Finally, the adaptability of this approach is a major benefit. Using a GFP nanobody linked to the TurboID enzyme makes this system modular and adaptable to a wide array of bait proteins, given the large number of GFP-tagged proteins in *C. elegans*. All that is required is a simple cross to introduce both components into a single worm strain. Further, the selection of tissue and cell-type specific promoters to drive TurboID expression enables a high degree of specificity in labeling. Altogether, this modular approach to proximity labeling has significant potential for uncovering new aspects of protein dynamics and function *in vivo*, which is beginning to play out in the literature.

### Overlapping datasets yields insights into potential germ granule components and pathways

By overlapping our TurboID datasets from eight AGOs, 2 RdRPs, and ZNFX-1 with existing TurboID data on GLH-1 and DEPS-1, we were able to define a high confidence set of proteins likely to be associated with germ granules (Fig 3A). Some of these factors are known, *bona fide* germ granule factors, while others remain to be fully studied for their role in germ granule biology. We characterized eleven of these factors, which we had prioritized based on their interaction with three or more baits. Other labs have also characterized some of these factors to varying extents, and our data are mostly consistent with these studies. Ultimately, much more work remains to be done to understand the full repertoire of molecular roles of these factors in germ granule biology, including full mutational analysis (including structure-function analysis) via CRISPR-Cas9 genome editing and TurboID on each of these proteins, as we did here for D2005.4, to identify proximal protein interactors. In addition to this set of proteins that interact with many of our baits, we uncovered many other uncharacterized factors that interact with only one or two baits that warrant further investigation (Fig 3A and S6).

Another category of proximal interactors that deserves further study are metabolic pathway proteins and enzymes resident in the ER, cytoplasm, and mitochondria, which were present in most data sets and, interestingly, different AGOs enriched for different subsets of these proteins. While many of these are abundant proteins that could be artifacts of this approach, the fact that we observed different metabolic factors associated with different baits may argue against this. In fact, metabolic pathway factors have been implicated in RNA binding, moonlighting as non-sequence specific RBPs, so it is plausible that these proteins could operate in conjunction with sRNA pathways and germ granules. In addition, many *ago* null mutants do not have phenotypes in the normal laboratory environment, and phenotypes are only revealed under stressful conditions^3^. This points to a major role for AGOs in buffering stress pathways. Because germ granules, particularly Z granules, are involved in epigenetic inheritance, it may be that by linking metabolic pathways and stress responsive pathways in the endoplasmic reticulum and mitochondria to germ granule physiology and sRNA mediated gene regulation, animals can respond to and “remember” environmental inputs or stresses for many generations. Sensing a stressor such as low oxygen, temperature, pathogens, etc. and linking the responses to germ granules and sRNA regulation could ensure survival in a single generation and potentially an adaptive heritable response for future generations. Further investigation will be necessary to determine the role–if any–for such metabolic and stress-related factors in germ granule physiology and epigenetic inheritance. For instance, determining if these proteins localize to germ granules, and identifying any transgenerational epigenetic inheritance phenotypes (e.g., RNAi inheritance, Mrt, etc.) would be a good starting point.

### D2005.4: a scaffold for sRNA pathways and link between germ granules and mitochondria?

D2005.4 is exceptional because it was identified as a proximal interactor with all AGOs. Yet, the converse-AGO association with D2005.4-was not observed. This was curious, but could be explained by several possibilities, including that D2005.4 is GFP tagged at the C terminus of the protein within an IDR, and if the interaction between D2005.4 with AGOs is within the TPR region or the N terminal IDR, it may be beyond the maximum distance of proximity labeling. The fact that we did not observe the AGOs in the D2005.4 TurboID network could also be reflected by the function of D2005.4. Because this protein has a central TPR region, we hypothesize that it could serve as a scaffold or chaperone for molecular assemblies of many proteins involved in sRNA processing of 22G-RNAs and piRNAs, along with post-transcriptional regulation. The protein also possesses N and C terminal IDRs. IDRs are often associated with phase separation, however our deletion analysis demonstrated that D2005.4’s IDRs were not required for germ granule localization, leaving the TPR domain, or interactions with other proteins or even RNAs as a means for recruitment to condensates. The enrichment of mitochondrial proteins in the D2005.4 interaction network was notable, and may highlight an under-appreciated linkage between germ granules and mitochondria. While a preliminary experiment to look for D2005.4 co-localization with mitochondria did not show overwhelming overlap (data not shown), this point should be revisited with live imaging experiments, as we anticipate the interaction of D2005.4 with mitochondria may be transient. In other organisms such as *Drosophila* and mammals, germ granules and mitochondria are in physical proximity within germ cells and are functionally linked^109,110^. In fact, the mitochondria provides a substrate for the synthesis and processing of piRNAs via the ping pong cycle^111^. piRNAs are generated by a different process altogether in *C. elegans*, and the amplification occurs via 22G-RNAs, rather than piRNAs themselves, in germ granules. Nevertheless, the abundance of mitochondrial factors in our TurboID indicates that this relationship should be studied further.

### Identifying proximal interactors has the potential to shine a light on AGO and RNAi pathway factor functions and dynamics

Despite having been studied for many years, the exact functions of AGOs in gene regulation remain mostly elusive. For AGOs without endonucleolytic activity, regulatory mechanisms appear to be restricted to sequestering target RNAs, dismantling RNAs, recruiting nucleases to degrade RNAs, inhibiting ribosome association and translation initiation, and perturbing translation^4^. Nuclear AGOs are generally thought to influence chromatin, or modulate RNA Polymerase activity^112^. Our TurboID data have provided new insights about potential mechanisms of action for *C. elegans* AGOs and highlighted unknown aspects of the dynamic cycle of AGOs within the germline.

WAGO-4 provides a clear example of our TurboID data shining a light both on the localization and function of AGOs. WAGO-4 has been thought to be restricted to the cytoplasm and germ granules. However, observing WAGO-4 with proximal interactors involved with chromatin and the kinetochore in the 3xFG background led us to explore the possible nuclear localization of WAGO-4. Treatment of GFP::3xFLAG::WAGO-4 worms with the nuclear export inhibitor, leptomycin B, revealed a pool of nuclear WAGO-4 in both backgrounds (Fig 2D). Upon further inspection, this pool was visible in the 3xFG background even without leptomycin B treatment. This was a surprising finding that explains the interaction of WAGO-4 with chromatin and nuclear factors in the 3xFG background. We speculate that we only detect these chromosome proteins in the 3xFG background because in WT worms, WAGO-4 is concentrated in germ granules, which clearly predominates the interactors we recover in our TurboID data (Fig 2A). While WT WAGO-4 displayed an association with factors involved in germ granules, like helicases, and post-transcriptional regulation of mRNAs, the number of helicases associated with WAGO-4 in the 3xFG background decreased and the number of RNA regulatory protein proximal interactors increased. A nuclear role for WAGO-4 has not been observed prior to this. However, the 3xFG mutant results in an endomitotic oocyte phenotype at 26 °C, which we speculate is related to this newfound nuclear pool of WAGO-4^25^.

WAGO-4 lacks the residues required for endonucleolytic activity, therefore it was expected that it would act post-transcriptionally and via co-factor recruitment. Indeed, mRNA-seq experiments have shown an increase in WAGO-4 target transcript levels upon *wago-4* mutation, indicating a destabilizing action on targets^24^. In our TurboID data, we recovered proximal interactors involved in both 5′ and 3′ RNA regulation, including scaffolds for de-capping and de-capping enzymes, de-adenylases, RNA binding proteins, and exonucleases, pointing to WAGO-4 as a key post-transcriptional regulator involved in RNA decay. That these interactions were further enriched in the 3xFG background points to the cytoplasm as a key point of post-transcriptional regulation by WAGO-4. Taking this observation one step further, it was the set of CSR-1 shared targets that were repressed in the 3xFG background, while a smaller pool of WAGO-4 targets not shared with CSR-1 were de-repressed, indicating that the shared WAGO-4/CSR-1 targets are likely to be subject to WAGO-4 mediated RNA decay in the cytoplasm, while the WAGO-4 specific targets may be regulated by similar mechanisms, but in the germ granules.

Another AGO that provided a surprise was the nuclear AGO HRDE-1 and its set of proximal interactors involved in splicing. While a large body of work has implicated HRDE-1 in the formation of heterochromatin at target gene loci, in only one study was HRDE-1 previously associated with the intron binding protein, Aquarius/EMB-4, which is involved in splicing^85^. Our TurboID data recovered no histone methyl-transferases, yet enriched for proteins involved in each stage of splicing (Fig 1C and S4). HRDE-1 is also enriched for proteins interacting with the C-terminal domain of RNA Polymerase II, and for components of the Integrator complex. The Integrator complex is involved in snRNA and piRNA transcriptional termination, and in the termination of some protein coding genes. Notably, recent studies using acute degradation of HRDE-1, rather than prolonged absence of the protein, demonstrate no associated alterations in heterochromatin as an acute effect of *hrde-1* loss, pointing to this as a likely indirect or secondary outcome of the HRDE-1 pathway^73^. What role HRDE-1 plays in splicing regulation is not yet clear.

CSR-1 is the only essential AGO, and the only one that resides in a distinct position within the germ granules, at the interface between the nuclear pore and P Granule as a part of the D Granule/Compartment^13^. This role is clearly evidenced by being the only AGO associated with nuclear pore complex components and DDX-19. Because CSR-1 is required for the licensing and protection of nearly 5000 germline expressed protein coding genes^3^, we speculate it may be part of a scanning and sorting mechanism for transcripts emerging from the nucleus, to be routed to appropriate locations in germ granules or the cytoplasm. In addition, the fact that CID-1/CDE-1 was identified as an interactor of all AGOs but HRDE-1, yet showed the most plausible model with CSR-1 indicates its key role in uridylating CSR-1 22G-RNAs, as was initially hypothesized when CID-1/CDE-1 was first linked to CSR-1^39^ (Fig 1D). Instead of uridylating the 22G-RNAs bound to other AGOs, CID-1/CDE-1 may assist in handing off 22G-RNAs that are uridylated to other AGOs, or may be part of the 22G-RNA loading/unloading pathway in some other capacity.

WAGO-1 and PPW-2 protein interaction networks, particularly that of WAGO-1, showed the most overlap of any AGOs with each other and with PRG-1, reflecting the connection between the primary piRNA pathway and the biogenesis of 22G-RNAs (Fig 1C). The proximal interaction of all three factors with PID-2 and PID-5 also highlights their known links to TEI mechanisms in association with Z granules. The interaction of all three factors with the eIF4E transporter, IFET-1 may point to a role in translational inhibition and storing transcripts in germ granules. IFET-1 interacts with the eIF4E (cap binding protein) paralog, IFE-3^113^. While its primary role is as a protein that transports IFE-3 between the nucleus and cytoplasm^114^, it can also act as a protein that blocks the interaction between eIF4E and eIF4G, prohibiting ribosome assembly and holding transcripts in a state of stasis^115^. We speculate that these AGOs may recruit or stabilize IFET-1 on transcripts that are stored, poised for translation, in germ granules.

Our final surprise came from RRF-1. While a previous study hinted at a potential nuclear role for an RdRP^85^, our TurboID data emphasized nuclear proteins involved in nuclear transport, in the DSB pathway, and transcription termination (Fig 3A). Leptomycin B treatment identified a clear nuclear pool of RRF-1, and RNAi treatment for *ipo-13* demonstrated its role in the nuclear accumulation of RRF-1 (Fig 3B). Gamma irradiation to induce DSBs led to a decreased brood size for rrf-1 mutants, indicating they are more sensitive to DSBs and implicating them in this process (Fig 3C). In *Arabidopsis* and humans, sRNA pathways, including a specific subset of sRNAs, Dicer, AGO, and in *Arabidopsis*, a specific RNA Polymerase that generates sRNAs, have been implicated in DSB sensing and repair^116^. It is plausible that RRF-1 could synthesize sRNAs at DSB sites that may help to degrade transcripts and stop transcription at break sites, or that may help to signal the damage. While the involvement of Dicer in this process is unclear, it is tempting to speculate that a nuclear AGO such as HRDE-1 could play a role in this process, or, based on its TurboID interaction with RAD-50, that PPW-2 may play a role. Clearly, further work is required to clarify the mechanisms by which RRF-1 and any other RNAi factors could participate in the DSB pathway and other nuclear functions.

Collectively, these results highlight the utility of TurboID as a tool to develop new hypotheses and gain new insights into germ granule and sRNA component functions. Our work also reminds us that even after decades of study, there are still many mysteries of sRNA pathway and germ granule function to unravel, and that these proteins are not finished with surprising–and delighting–us.

## METHODS

### Nematode maintenance and strain generation

Unless otherwise stated, all strains used in the study were grown at 20 °C on 3.5 cm Nematode Growth Medium (NGM) plates inoculated with *Escherichia coli OP50* bacteria^117^. GFP::3xFLAG tagged strains were previously generated via CRISPR-Cas9 mediated genome editing^3^.“Indirect” TurboID strains were made via crossing GFP tagged strains with the TurboID-anti-GFP nanobody fusion strain (DAM1284) provided by the Dammerman Lab^33^. A complete list of strains used in this study are provided in Supplemental Table S1. Primer sequences for genotyping are found in Supplemental Table S2. The Bristol N2 strain was used as a reference strain.

### TurboID sample preparation

Strains used for TurboID were synchronized, grown, and harvested as previously published^34^. Flash frozen pellets were mixed with an equal volume of modified RIPA buffer (50 mM Tris-HCl, 150 mM NaCl, 0.1 % SDS, 0.5 % sodium deoxycholate, 1 % Triton X-100, 1 mM EDTA, 2 mM DTT, Roche Protease complete inhibitor 2 tablets (Sigma), 0.1 % phosphatase inhibitor-2 (Sigma), 0.1 % phosphatase inhibitor-3 (Sigma) and manually homogenized with a pestle. Homogenized worm lysates were sonicated and treated with 250 units/ml Turbonuclease for 15min. Turbonuclease was inactivated by increasing the SDS concentration of the samples to 0.4 %. Lysates were centrifuged at 13,500 x *g* for 15 min at 4 °C and the soluble lysate was transferred to a fresh tube. To determine total protein concentration, a Lowry assay was performed using a NanoDrop 1800C spectrophotometer (Thermo Fisher).

### Affinity purification

Pre-clearing was performed using agarose beads (ThermoFisher) and rotated at 4 °C for 1 hr. Samples were then centrifuged at 500 x *g* for 1 min at 4 °C and the supernatant was transferred to a fresh tube where it was combined with 40 µl 50 % v/v streptavidin-conjugated sepharose beads (Sigma) triple-washed with RIPA buffer and rotated at 4 °C for 1 hr. Following centrifugation at 500 x *g* for 1 min at 4 °C, supernatant was transferred to a fresh tube and stored for downstream analysis while beads were processed for streptavidin blots or mass spectrometry analysis as described below.

### Mass spectrometry preparation

Sepharose beads from the previous section were washed at 4 °C for 7 x 10 min with DROSO buffer (30 mM Hepes, 100 mM Potassium Acetate, 2mM Magnesium Acetate). Samples were then submitted to the Network Biology Collaborative Centre (NBCC) at the Lunenfeld-Tanenbaum Research Institute where they underwent on-bead trypsin digestion followed by nano-LCMS/MS mass spectrometry in Data Dependent Acquisition (DDA) mode, as detailed below.

### TripleTOF6600 acquisition

One quarter of each sample was acquired on a ABSciex TripleTOF™6600 in data-dependent acquisition (DDA) mode. Digested peptides were analyzed using an Eksigent ekspert™ nanoLC 425 coupled to a TripleTOF™6600 mass spectrometer. Peptides were separated on a home-packed column (100 µm internal diameter, 365 µm outer diameter and 5-8 µm emitter opening, packed with C18 reversed-phase material (Reprosil-Pur 120 C18-AQ, 3 µm)), with an emitter generated by a laser puller (Sutter Instrument Co., model P-2000, with parameters set as heat: 280, FIL = 0, VEL = 18, DEL = 2000). Sample in 5 % formic acid was directly loaded at 800 nL/min for 20 minutes onto the column and eluted with a linear 90-minute gradient from 2 % acetonitrile with 0.1 % formic acid to 35 % acetonitrile with 0.1 % formic acid. The gradient was followed by a 15-minute wash of 80 % acetonitrile with 0.1 % formic acid. After the wash, the column was equilibrated for 15 minutes with 2 % acetonitrile with 0.1 % formic acid in preparation for the next sample. The overall length of the DDA method was 135 minutes. The MS1 scan had a mass range of 400-1800 Da, with an accumulation time of 250 ms. This was followed by 10 MS2 scans of the top 10 most abundant precursors identified in the MS1 scan. Filters for each MS2 scan included the candidate ion to have a charge state from 2-5, have a minimum threshold of 300 counts per second, and previously analyzed candidate ions were dynamically excluded for 7 seconds. Each candidate ion was isolated using a window of 50 mDa and was allotted a maximum accumulation time of 100 ms.

### timsTOF Pro 2 acquisition

One sixteenth of each sample was acquired on a Bruker timsTOF Pro 2 in data-dependent acquisition (DDA) mode. Digested peptides were analyzed using an Evosep One coupled to a timsTOF Pro 2 mass spectrometer. The Evosep One was coupled to the timsTOF Pro 2 using a 20 µm diameter emitter tip (Bruker). The column toaster was set to 40 °C. Peptides were separated on a Performance column (Evosep, Cat#: EV-1109, 8 cm x 150 µm, packed with 1.5 µm beads). The overall length of the DDA method was 22 minutes. The MS1 scan had a mass range of 100-1700 Da in PASEF® mode, within the mobility range (1/K0) of 0.85 to 1.3 V·s/cm2. 1+ ions were excluded from fragmentation using a polygonal filter. Both accumulation and ramp time were set to 100 ms (with 4 PASEF ramps and active exclusion at 0.4 min). The target ion intensity was set to 17,500 with intensity threshold at 1750. The cycle time was 0.53 seconds.

### Data search and analysis

TripleTOF™6600 DDA and timsTOF Pro 2 data was stored, searched, and analyzed using ProHits laboratory information management system (LIMS) platform. For TripleTOF™6600 data, WIFF files were converted to an MGF format using the WIFF2MGF converter and to an mzML format using ProteoWizard (V3.0.10702) and the AB SCIEX MS Data Converter (V1.3 beta). The data was then searched using Mascot (V2.3.02) and Comet (V2018.01 rev.4). The spectra were searched with the *Caenorhabditis elegans* sequences in the RefSeq database (*Caenorhabditis elegans*, version 73, December 18, 2015) acquired from NCBI, supplemented with “common contaminants” from the Max Planck Institute (http://www.coxdocs.org/doku.php?id=maxquant:start_downloads.htm) and the Global Proteome Machine (GPM; ftp://ftp.thegpm.org/fasta/cRAP) and forward and reverse sequences (labeled “gi|9999” or “DECOY”) for a total of 56,564 entries. Database parameters were set to search for tryptic cleavages, allowing up to 2 missed cleavage sites per peptide with a mass tolerance of 35 ppm for precursors with charges of 2+ to 4+ and a tolerance of 0.15 amu for fragment ions. The search included variable modifications of deamidated asparagine and glutamine and oxidized methionine. ^118^Results from each search engine were analyzed through TPP^119^ (the Trans-Proteomic Pipeline, v.4.7 POLAR VORTEX rev 1) via the iProphet pipeline^120^.

For timsTOF Pro 2 DDA data, files were searched with MSFragger v.4.1 within the ProHits LIMS. The spectra were searched with the *Caenorhabditis elegans* Uniprot proteome, UP000001940, with no isoforms. Acetylated protein N-term and oxidated methionine were set as variable modifications. Precursor mass tolerance was set to 20 ppm on either side. Fragment mass tolerance was set to 20 ppm. Enzymatic cleavage was set to trypsin with 2 missed cleavages. MSBooster and Percolator were turned on. Percolator required a minimum probability of 0.5 and did not remove redundant peptides. The target-decoy competition method was used to assign q-values and PEPs. For ProteinProphet, the maximum peptide mass difference was set to 30 ppm. When generating the final report, the protein FDR filter was set to 0.01. FDR was estimated by using both filtered PSM and protein lists. Razor peptides were used for protein FDR scoring. All other parameters were default.

To determine high-confidence proximal protein interactors, samples for TripleTOF™6600 and timsTOF Pro 2 were analyzed separately with SAINTexpress v.3.6.3^121^. For TripleTOF™6600 data, protein results were filtered by TPP Probability >= 0.95 and required at least 2 unique peptides prior to SAINTexpress analysis. For timsTOF Pro 2 data, proteins needed at least 2 unique peptides to be included in the SAINTexpress analysis. For each analysis, N2 and DAM samples were used as negative controls. Controls were compressed to n = 10 with no bait compression^36–38,122^. Unique preys or those with 1.5x greater average spectra vs nanobody control (DAM1284) with a BFDR ≤ 0.05 (scored against N2) were identified as significant hits. Protein interaction maps (dot plots) were created with custom R scripts, available at GitHub (https://github.com/jmclaycomb/TurboID_AGOs). Manual curation and assignment of proteins to various groups was performed, using information from various sources, including Wormbase and the Alliance of Genome Resources, the STRING database, UniProt, and g:Profiler^123–127^.

### Streptavidin and western blots

Streptavidin blots were performed using previously established protocols with a few notable exceptions^3,34^. Sepharose beads from TurboID sample preparation were boiled in 1X Bolt™ Reducing Agent (ThermoFisher), 1X Bolt™ LDS Sample Buffer (ThermoFisher), and 3 mM biotin at 95 °C for 15 min. 0.5 % of the input/lysate, 1 % of the supernatant after pulldown, and ⅓ of the pulldown was loaded onto 10 % SDS-PAGE gels (Tris-Glycine). Membranes were incubated with a 1:2000 dilution of streptavidin-HRP (1.25mg/ml) (Thermo Scientific). Luminata Forte Western HRP substrate (Millipore-Sigma). Dilution series western blots for D2005.4::GFP::3xFLAG (WT) and the IDR C and N terminal deletion mutants consisted of 25 μl of lysate equating to 100 worms, 50 worms, and 25 worms for each strain. Western staining and imaging were performed using anti-FLAG (1:1000), and anti-Tubulin (1:10000) antibodies and protocols consistent with previous work in our lab^34^.

### Leptomycin B treatment

Leptomycin B (LMB; BioShop) was diluted in *OP50* bacterial culture and seeded onto NGM plates to a final concentration of 750 ng/mL. For vehicle control, an equivalent volume of absolute ethanol was used. Synchronized late L4-stage worms (52 h post-L1 arrest) were transferred to LMB or control vehicle plates and incubated at 20 °C for 4 h protected from light prior to imaging.

For T16G12.6/IMPORTIN 13 RNAi experiments, LMB or ethanol as vehicle control was diluted in T16G12.6 RNAi or L4440 bacterial culture, induced to express dsRNA with 5 mM IPTG at 37 °C for 4 hr, and seeded onto NGM plates to a final concentration of 750 ng/mL.

### Germline microscopy and image analysis

Twenty-five to forty adult hermaphrodites were mounted on 2 % agarose pads on slides and immobilized with 20 mM NaN_3_ or 2mM levamisole. Full germlines or only the pachytene region of the germline was imaged at Leica SP8 inverted scanning confocal microscope using 63x (NA 1.4, immersion oil) objectives. All images presented are a single 0.3 μm slice at digital zoom of 1x, 2x, or 4x. All image processing and analysis was performed using FIJI/ImageJ^128^.

For determining germ granule or nuclear to cytoplasmic fluorescent ratios of GFP::3xFLAG AGO localization (Fig. 1A), strains JMC164, JMC213, JMC219, JMC221, JMC223, and JMC231 were captured as described above using 20 mM NaN_3_ with an additional 2x digital zoom to include a minimum of 20 nuclei per worm. For analysis, a custom pipeline was used whereby masks were created using a manually chosen threshold per channel, granule size, count, and AGO fluorescence were measured using the Analyze Particles plugin. The measurements from these analyses were processed and visualized in R^129^. All code used is available on GitHub (https://github.com/jmclaycomb/TurboID_AGOs).

For live imaging of GFP::3xFLAG::WAGO-4 in WT and 3xFG backgrounds after LMB/vehicle control treatments (Fig. 2D), worms were picked into a 1x M9 droplet with 2 μl of 2 mM levamisole on standard microscopy slides with 2 % agarose pads. All images were acquired with a Leica SP8 inverted scanning confocal microscope as above. For fixed imaging of GFP::RRF-1 after LMB/vehicle control treatments, germlines were excised on positively charged slides in EBT (1x Egg Buffer, 0.1 % Tween-20) frozen and cracked on dry ice for 10 min and fixed at -20 °C in methanol, 1:1 methanol-acetone, acetone, for 5 min each. Slides were washed 3x in PBST (1x PBS, 0.1 % Tween-20) for 10 min, 3x in PBS for 5 min, stained with DAPI for 10 min, followed by 3x PBS washes for 5 min at room temp. Samples were mounted in VECTASHIELD Antifade Mounting Medium (BioLynx, VECTH1000). All images presented are a single 0.3μm slice at zoom of 3x.

For colocalization analysis of RFP::ZNFX-1, PGL-1::BFP (Fig. 4C), RNAi against uncharacterized germ granule factors or empty vector L4440 was performed by feeding synchronized strains JMC260 and JMC306. TetraSpeck™ 0.1 µm microspheres were included on each slide to control for imaging artifacts, and images were captured at an additional 6x digital zoom to get a higher per-nucleus resolution. The MultiStackReg plugin^130^ was used to align the fluorescent channels based on the TetraSpeck microspheres, and three nuclei per worm were manually selected for analysis. The BIOP JACoP plugin^131^ was used to calculate the Manders Colocalization Coefficient using a manually selected threshold per channel.

For imaging biotin and bait proteins (Supplemental Fig. S2), dissected germlines were freeze cracked and subjected to methanol/acetone fixation as described previously^34^. Primary antiGFP antibodies (0.4mg/ml) (Milipore Sigma) were used at 1:100 dilution for GFP tagged EGO-1 and RRF-1 TurboID strains, anti-HA (100 μg/mL) (Milipore Sigma) was used for the direct ZNFX-1 TurboID strain, and anti-FLAG (1mg/ml) (Milipore Sigma) was used for remaining stains at 1:100 (Indirect ZNFX-1), 1:1000 (CSR-1), and 1:500 (all other stains). Secondary antibodies consisted of 1:500 dilutions of donkey anti-mouse FITC (FLAG stains) (Jackson immuno research), goat anti-mouse Alexa Fluor 488 (GFP stain) (Fisher scientific), and donkey anti-rat Alexa Fluor 488 (HA stain) (Jackson immuno research). All stained slides were additionally treated with 1:500 streptavidin Alexa Fluor 647 (2mg/ml) (Fisher scientific). Imaging was conducted as described above on the Leica SP8 confocal.

### Brood size, mortal germline, and Transgenerational GFP RNAi assays

RNAi by feeding was conducted as previously described against each of the uncharacterized factors.^132–134^ Brood size and mortal germline assays were conducted using described methods^3,34^. In mortal germline assays, worms were grown at a non-permissive temperature (25 °C). Each generation, starting with P0, four L4 worms per plate are picked and placed on a new plate. 10 plates per strain were propagated until they no longer produced offspring. Transgenerational epigenetic inheritance was assayed on strain SX1263 via RNAi against each of the uncharacterized Factors for one generation before and throughout the assay. Synchronized worms were fed with GFP or empty vector RNAi combined with the uncharacterized factor RNAi at a 1:1 ratio for one generation. Single F1 worms were transferred to individual plates after ensuring GFP knockdown. Subsequent generations were passaged using four randomly selected L4s. The remaining worms were washed off plates and a subset (∼40-100) worms/condition were placed in 96-well plates with 2mM levamisole and imaged Leica SP8 inverted scanning confocal microscope using 10x objective lens. Each well was imaged by stitching together 32 10x magnification images. to determine the level of GFP silencing. Images were blinded and counted, blinded counts were then analyzed and unblinded before being adjusted for internal controls and graphed. The duration of inheritance was measured as the generation in which at least 80 % of the worms regained GFP expression.

### Gamma irradiation and brood size

Synchronized young adult worms (56 hr post-L1 arrest) were exposed to 60 Gy γ-rays using a Caesium-137 source. Following irradiation, worms were transferred onto NGM plates, incubated at 20 °C and transferred to fresh plates every 24 hr until egg laying ceased, and hatched progeny were counted.

### AlphaFold3 modelling

*In silico* protein–protein interactions were predicted using AlphaFold v3.0.1. Amino acid sequences for each protein were retrieved from UniProtKB. For predictions involving sRNA, representative sRNA sequences were selected following differential expression analysis using sRNAseq-nf (https://github.com/imu93/sRNAseq-nf) with publicly available IP and input sRNA-seq libraries^3^.

Predictions were performed in two stages on high-performance computing clusters from the Digital Research Alliance of Canada. In the first stage, unpaired multiple-sequence alignments were generated using local AlphaFold databases, and each complex was modelled using two seeds (5 and 10) and five diffusion models. The resulting models and intermediate files were then used as input for the second stage, in which model inference was performed to generate the final complex predictions.

Protein Interface Visualization tool (PIVS) was used to assess the validity of the predicted interaction models. This tool is a pipeline of existing analyses^135–156^, and uses published thresholds to classify the plausibility, viability, and confidence of the predicted interaction. In all cases the highest ranked model was used for analysis.

### Phylogenetic analysis of uncharacterized proteins

To assess the evolutionary conservation of uncharacterized factors, we performed remote homology searches across 54 predicted nematode proteomes representing the five major nematode clades using Hidden Markov Models (HMMs). Candidate homologs were first identified using BLASTp (-evalue 1e-3, -qcov_hsp_perc 30, length ratio 0.7-1.3) and aligned with MAFFT. Protein-specific HMMs were built with HMMER (hmmbuild) and searched against the combined proteome database using hmmsearch. Hits were retained using stringent protein- and domain-level thresholds (full-sequence E-value ≤1×10⁻¹⁰, best-domain i-Evalue ≤1×10⁻⁵, domain score ≥30, HMM coverage ≥0.35 and bias/score ≤0.3).

We visualised the conservation patterns as a matrix of frequency alongside a maximum-likelihood species tree based on 2,625 BUSCO genes present in 70 % of the species using the BUSCO phylogenomics (https://github.com/jamiemcg/BUSCO_phylogenomics), and RaxML-ng (--model LG+I+G4+F --bs-metric fbp --bs-trees 100).

## SUPPLEMENTAL FIGURE LEGENDS

**S1.**
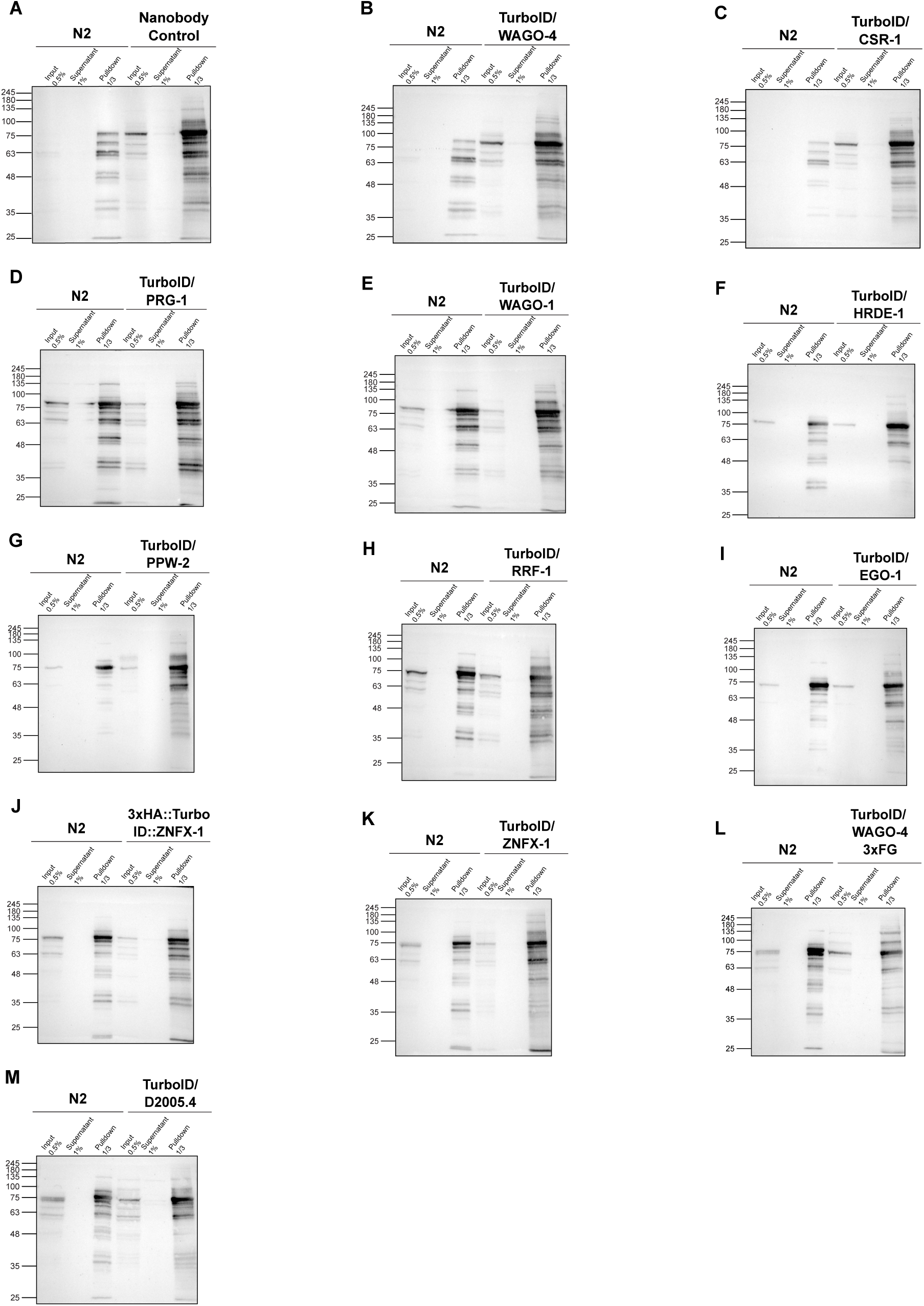
Streptavidin blots of bait strains and controls (related to Fig. 1) Streptavidin blots for each bait and TurboID only control, compared to N2 samples processed at the same time. Wells were loaded with equal volumes (15μl) diluted to the same concentration (3.33μg/μl). Percentage of total protein for each sample run is noted above each well (0.5 % of input, 1 % of supernatant, 33 % for pulldowns).

**S2.**
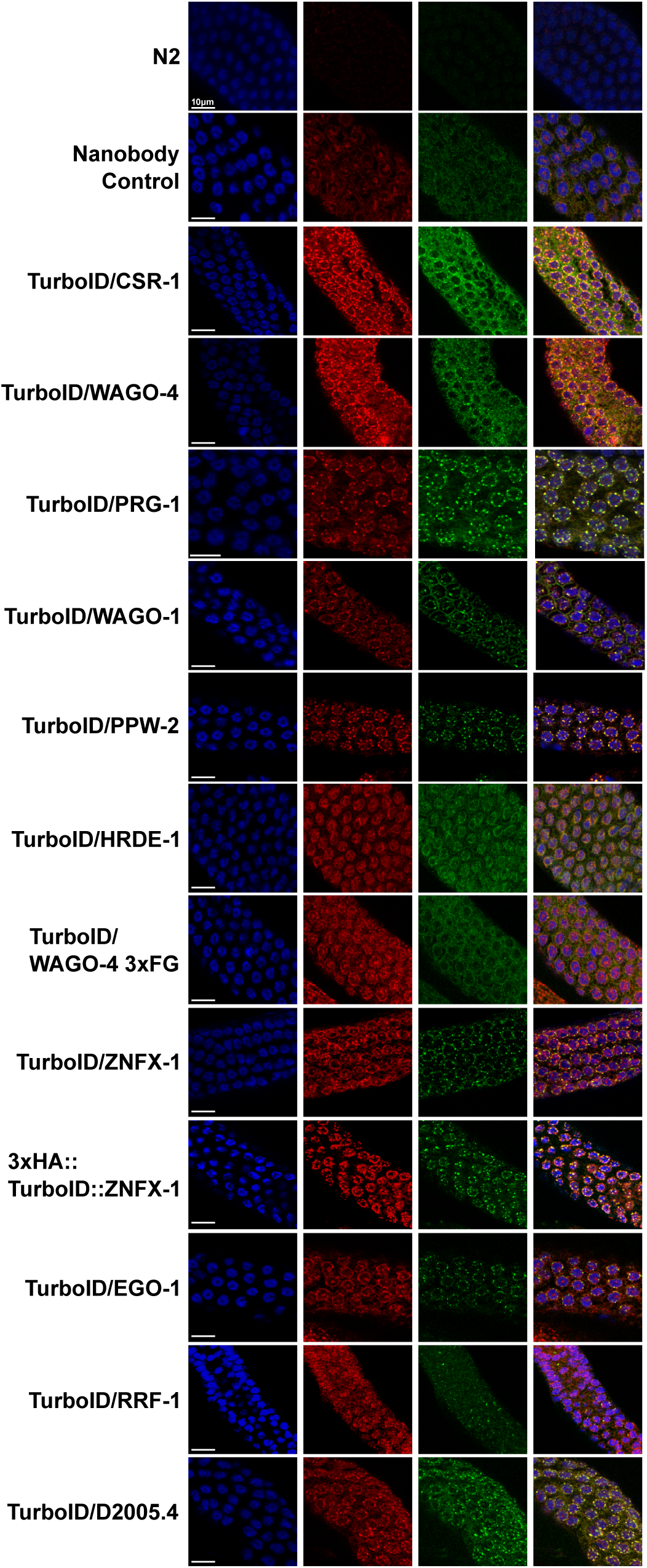
Streptavidin staining of bait TurboID strains and controls (related to Fig. 1) Stained dissected germlines for all bait TurboID strains used in the study. Anti-GFP (EGO-1, RRF-1), anti-HA (Direct TurboID ZNFX-1), and anti-FLAG (all other strains) staining was performed on the corresponding strains, alongside DAPI and streptavidin alexa fluor 647 staining. All images were taken using the same laser settings, LUTs were modified to highlight co-localization. Scale bar represents 10µm

**S3.**
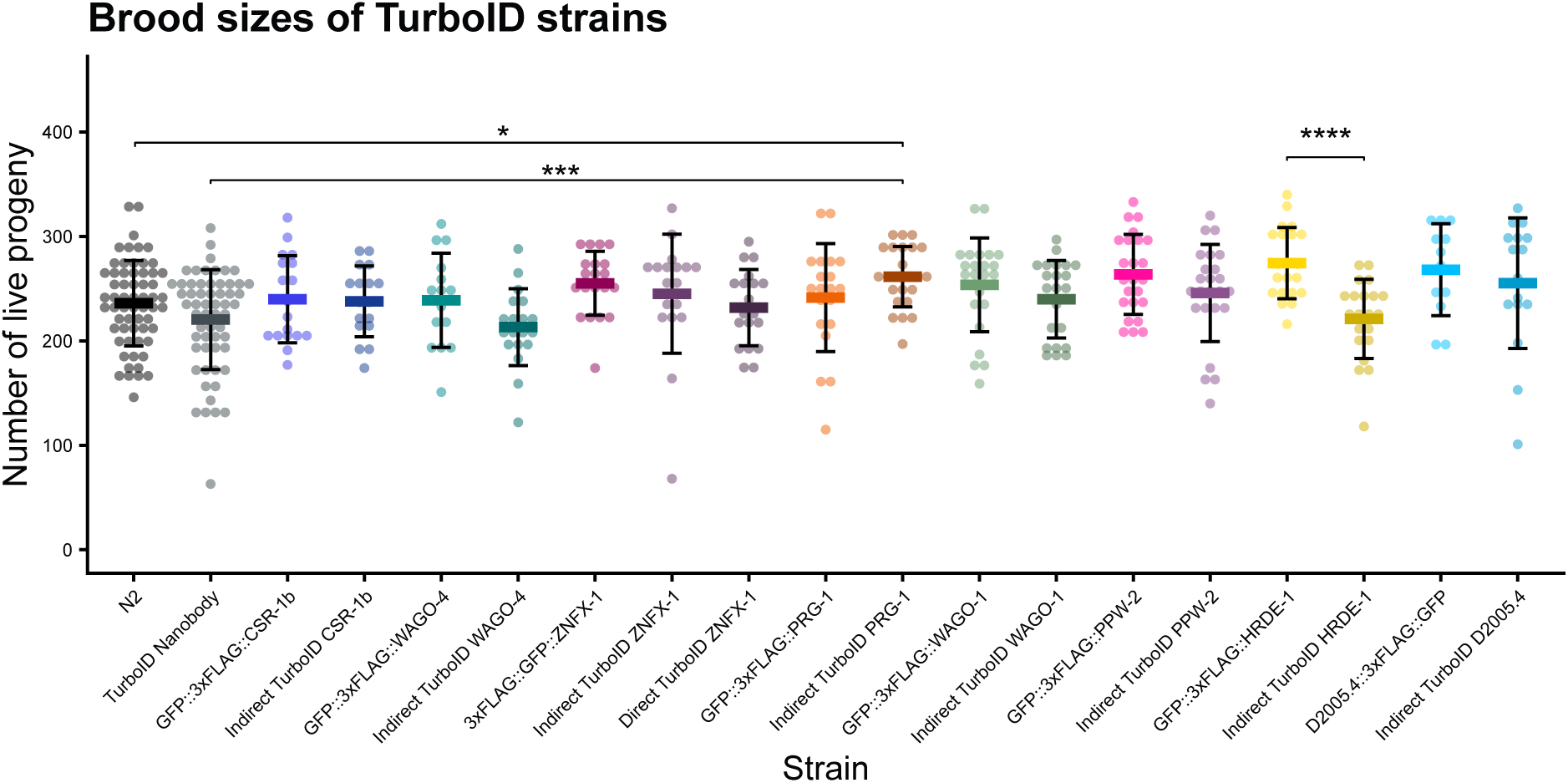
Brood size of bait strains (related to Fig. 1) Brood sizes of TurboID bait strains compared to TurboID only, WT (N2), and GFP tagged controls (without TurboID) at 20 °C. Note that various separate experiments were performed and compiled together, using the same controls. To correct for variation in the data, Dunnett’s t-test was used to compare bait TurboID strains against N2 and TurboID only strains, and a students T test was used to compare bait TurboID strains vs GFP controls (without TurboID) analyzed at the same time.

**S4.**
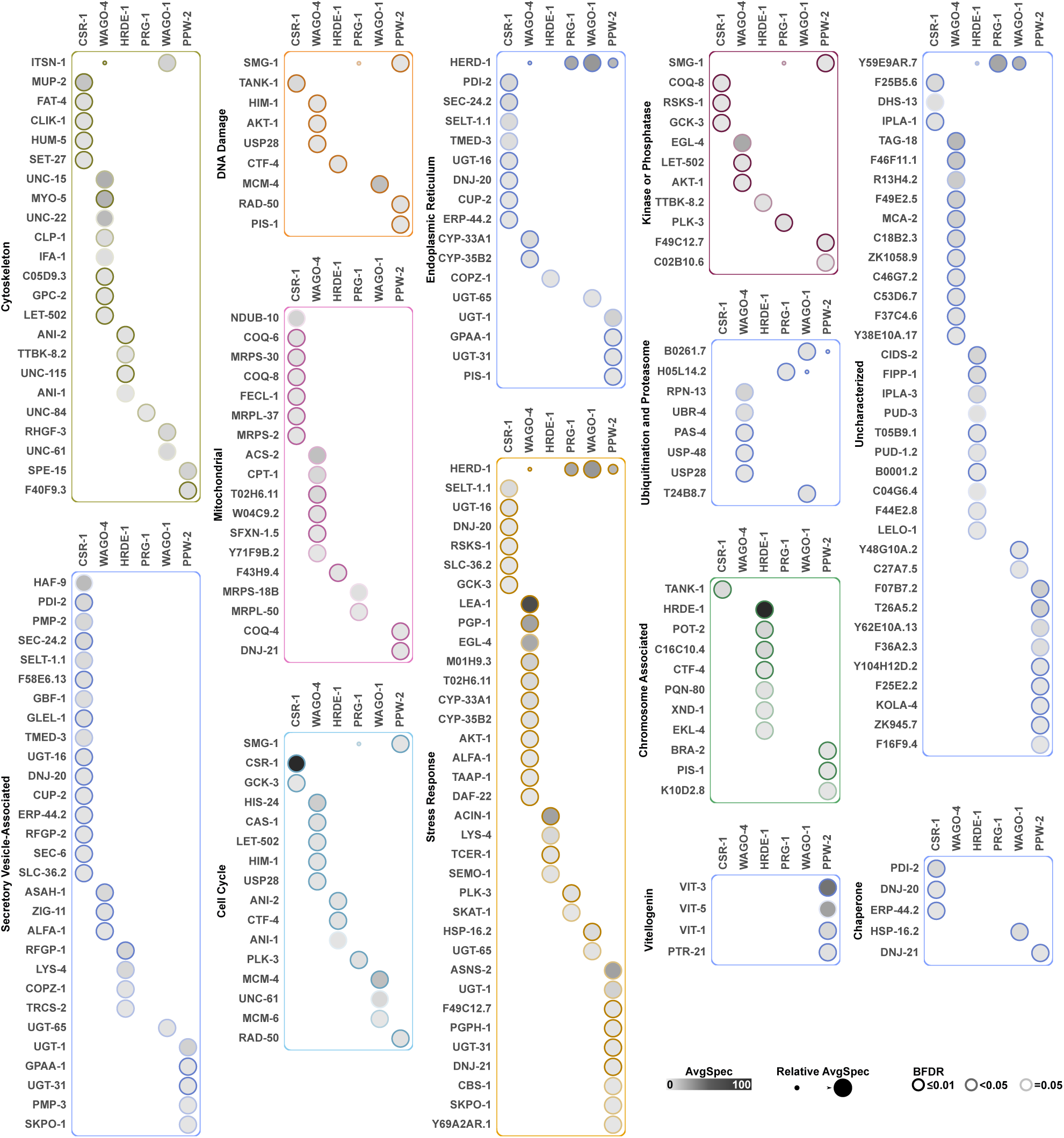
TurboID Dot Plots for AGOs (related to Fig. 1) Additional dot plots of AGO TurboID data, in which the average spectral count is represented by node colour intensity, and the Bayesian False Discovery Rate (BFDR) by solid (less than or equal to 0.01) or dashed (between 0.02 and 0.05) edges. Proteins are grouped by manually curated sets of proteins, and because of overlap in these categories, a proximal interactor may be observed in more than one category.

**S5.**
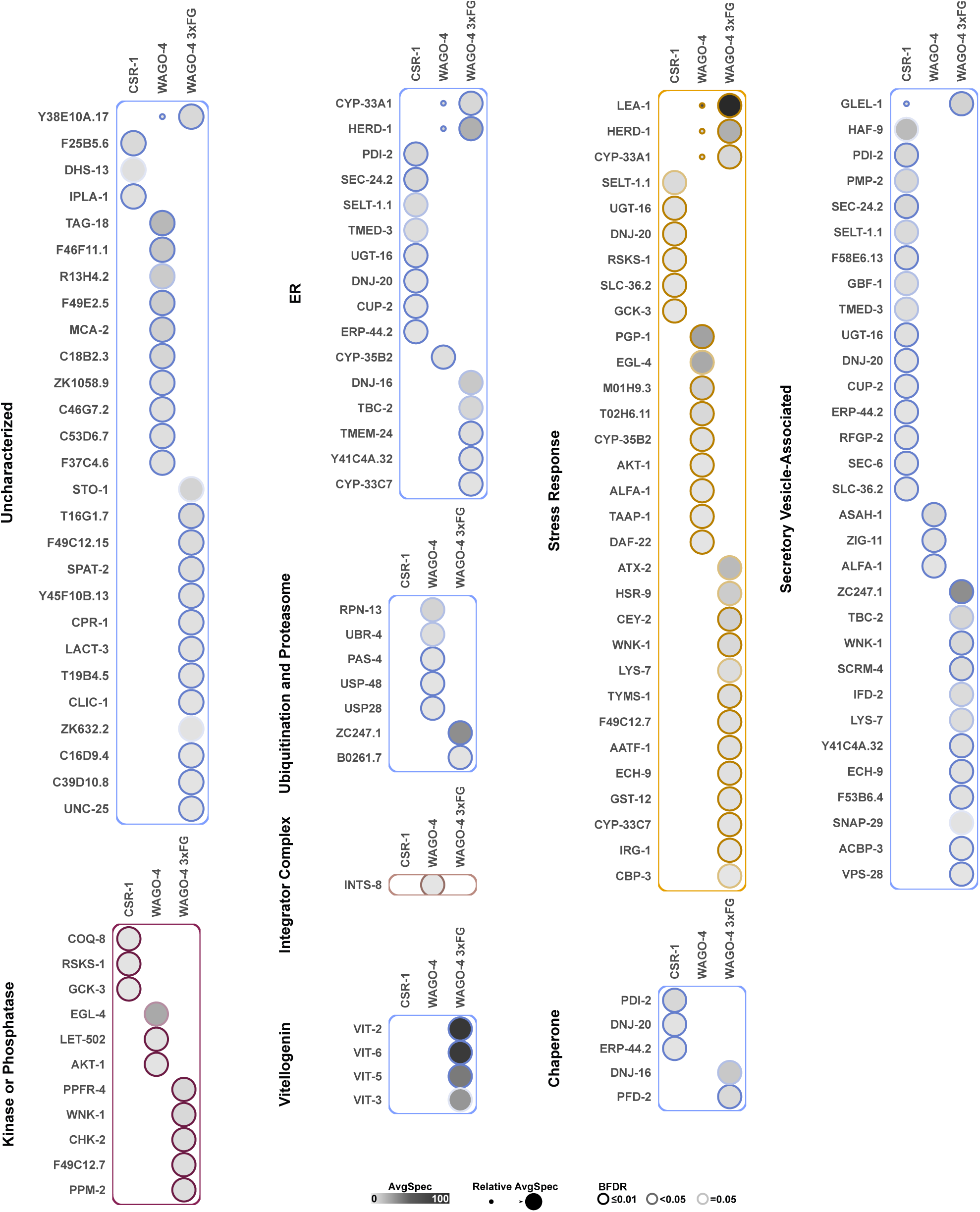
Turbo ID Dot Plots for WAGO-4 WT, WAGO-4 3xFG, and CSR-1 (related to Fig. 2) Additional dot plots of CSR-1, WAGO-4 WT, and WAGO-4 3xFG TurboID data, in which the average spectral count is represented by node colour intensity, and the Bayesian False Discovery Rate (BFDR) by solid (less than or equal to 0.01) or dashed (between 0.02 and 0.05) edges. Proteins are grouped by manually curated sets of proteins, and because of overlap in these categories, a proximal interactor may be observed in more than one category.

**S6.**
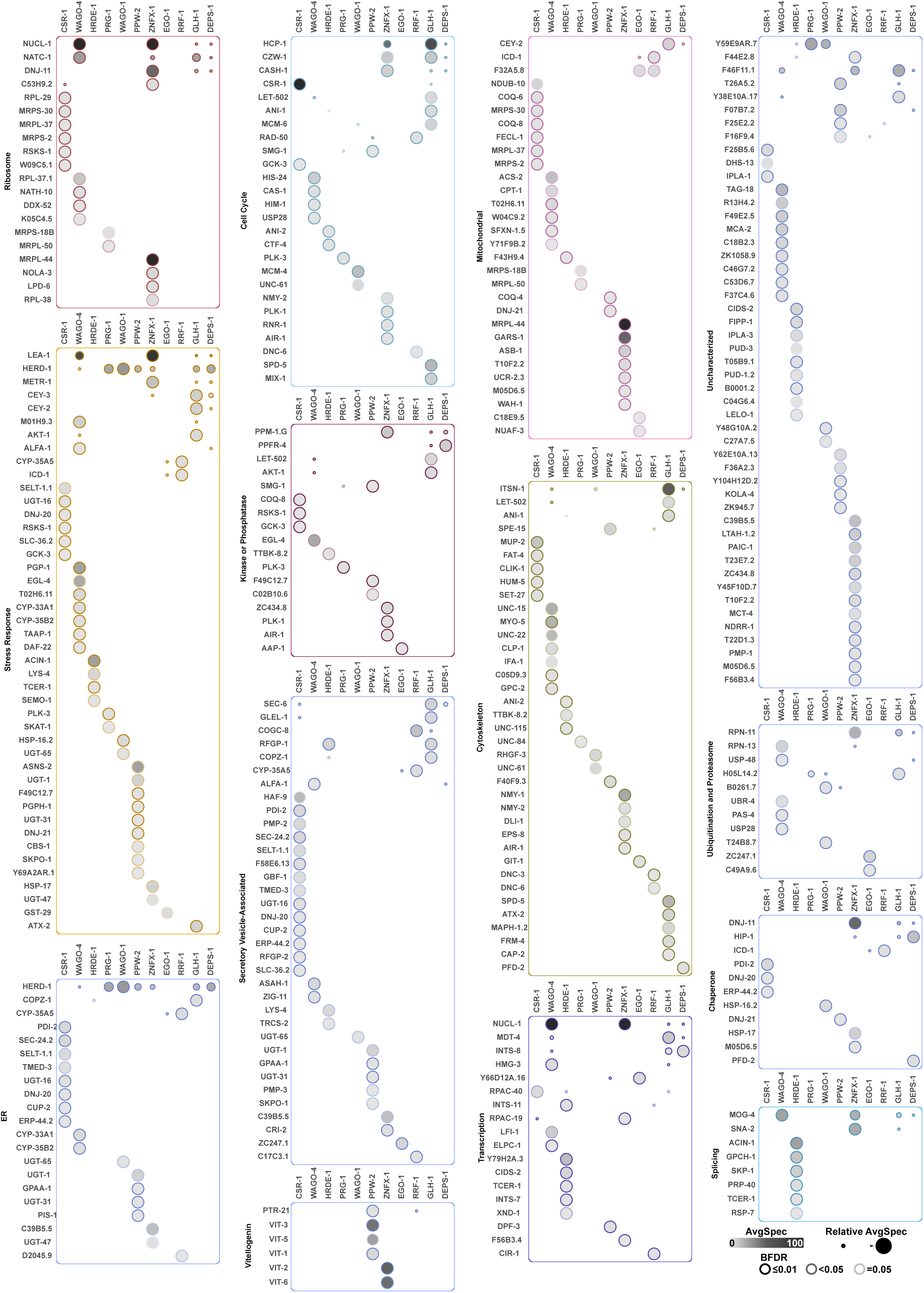
Turbo ID Dot Plots for all AGOs, ZNFX-1, RRF-1, EGO-1, GLH-1, and DEPS-1(related to Fig. 3) Additional dot plots of Fig. 3 TurboID data, in which the average spectral count is represented by node colour intensity, and the Bayesian False Discovery Rate (BFDR) by solid (less than or equal to 0.01) or dashed (between 0.02 and 0.05) edges. Proteins are grouped by manually curated sets of proteins, and because of overlap in these categories, a proximal interactor may be observed in more than one category.

**S7.**
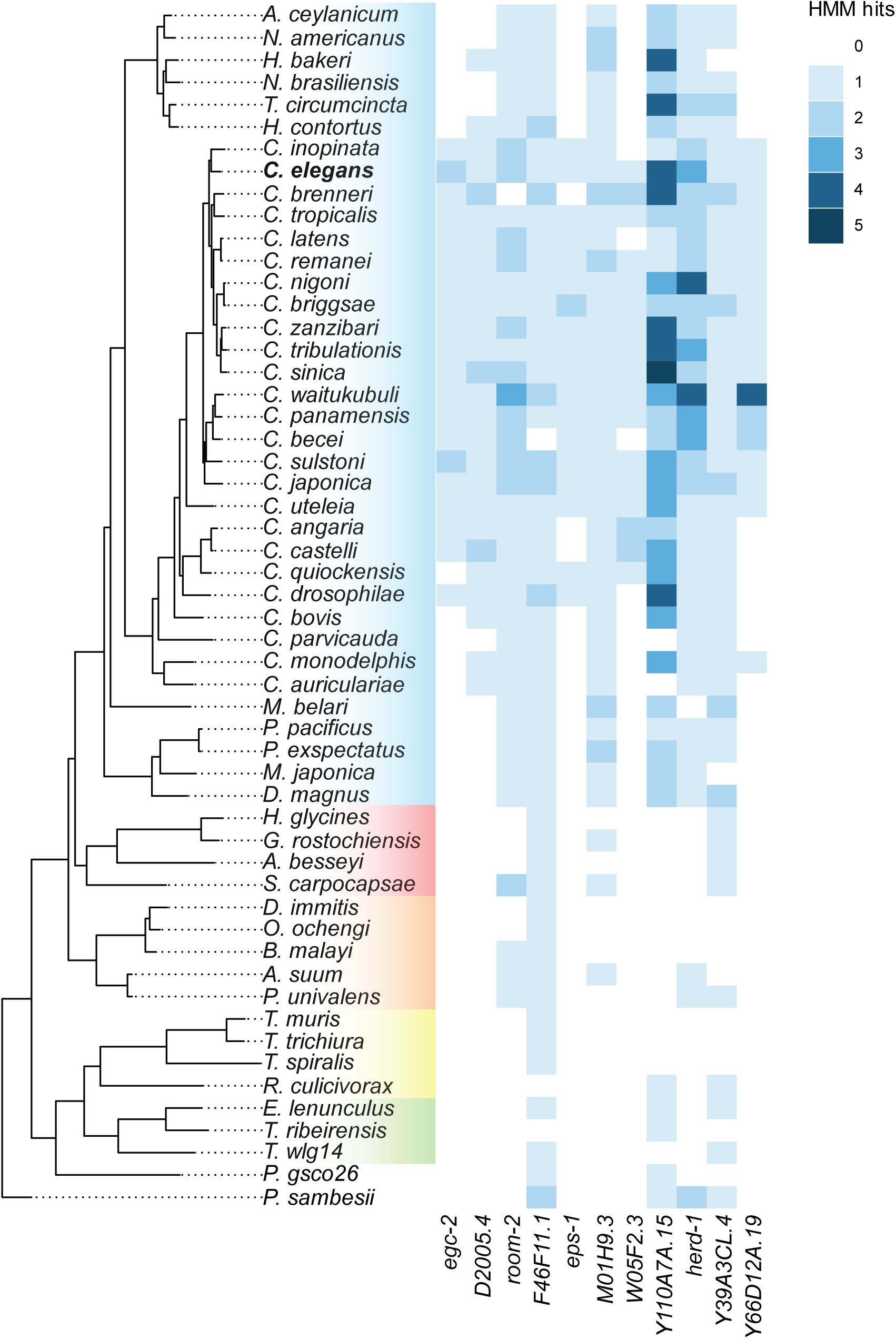
Phylogenetic tree of uncharacterized factors across 54 nematode species (related to Fig. 4)

**S8.**
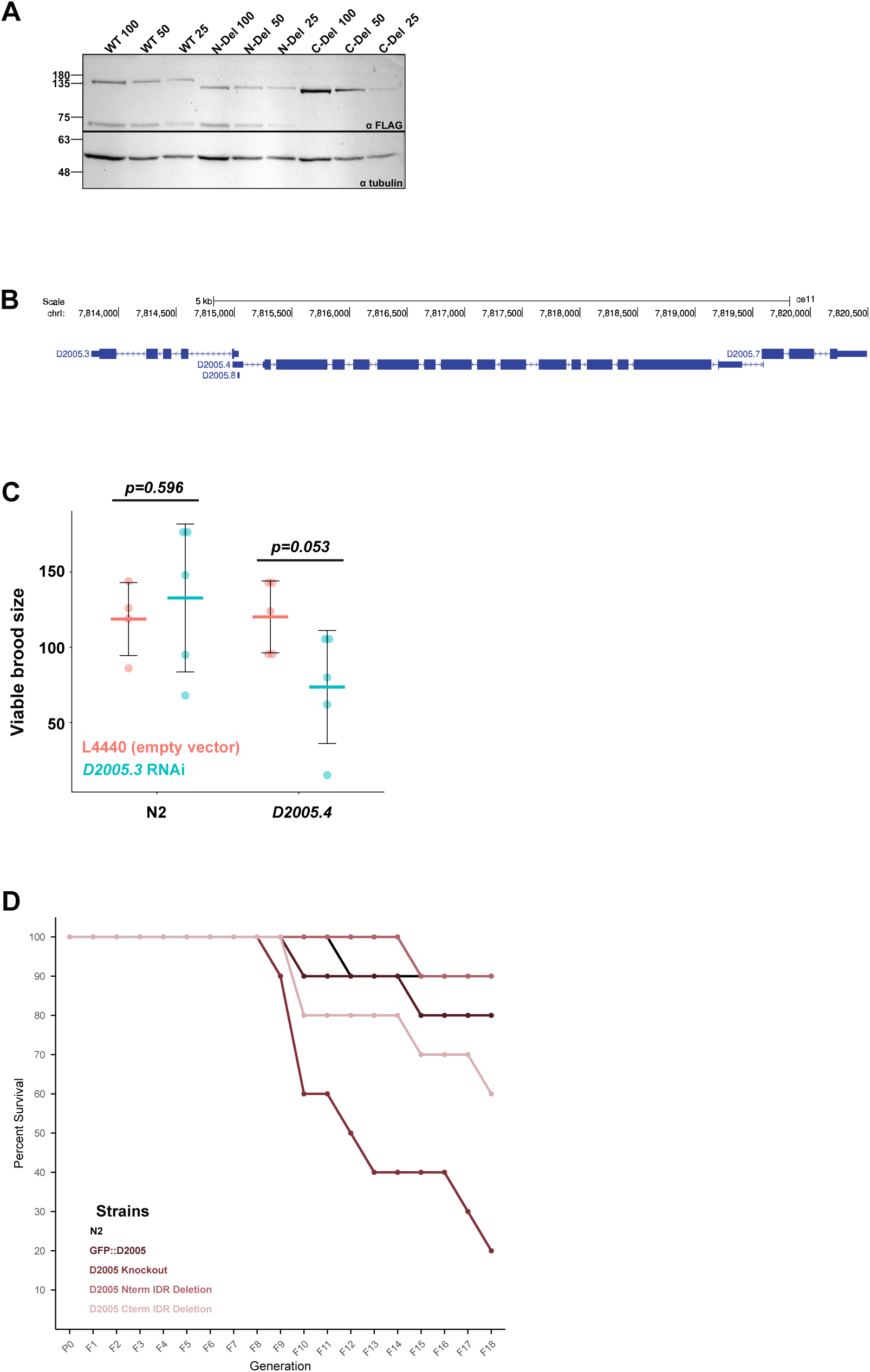
D2005.4 Supplemental Data (related to Fig. 5) **A.** Western blot used for quantification in Fig 5D. Dilution series for GFP tagged Wildtype D2005.4 (WT) and the two IDR domain mutants. Dilutions consisted of lysate equating to 100 worms, 50 worms, and 25 worms for each strain. Tubulin was used as a loading control. **B.** Genome browser diagram of the *D2005.3/.5/.7* locus, exported from UCSC Genome Browser **C.** RNAi of *D2005.3* in the *D2005.4* background. *D2005.4* deletion mutants were subjected to empty vector (L4440) or *D2005.3* RNAi from the L1 stage onward, and their viable brood at 25°C counted. *P* values are as noted with a students t test. N=5 worms per condition. **D.** Mortal germline assay of *D2005.4* mutants. Each strain was passaged at 25 °C for multiple generations, 4 worms were single picked to fresh plates each generation, with 10 plates/strain.

## SUPPLEMENTAL TABLES

**Supplemental Table S1: *C. elegans* strains used in this study**

**Supplemental Table S2: Primers used in this study**

**Supplemental Table S3: TurboID data from this study**

## DATA DEPOSITION

https://github.com/jmclaycomb/TurboID_AGOs All proteomic data are deposited at MassIVE.

## ACKNOWLEDGEMENTS

The authors wish to thank the Network Biology Collaborative Centre Proteomics Facility (RRID: SCR_025375) at the Lunenfeld-Tanenbaum Research Institute for mass spectrometry analysis. The facility is supported by the Canada Foundation for Innovation and the Ontario Ministry of Colleges, Universities, Research Excellence and Security. Some strains were provided by the CGC, which is funded by NIH Office of Research Infrastructure Programs (P40 OD010440).

We thank Adam Sundby, a former MSc student in the Claycomb lab who generated the TurboID::ZNFX-1 strain and started this project.

We thank the lab of Dr. Brent Derry (Sick Kids), in particular, Bin Yu for their assistance with the gamma-irradiation experiments, and we thank Dr. Michael Norris (University of Toronto Dept. of Biochemistry) for his expertise in examining protein-protein interactions.

Some strains were provided by the CGC, which is funded by NIH Office of Research Infrastructure Programs (P40 OD010440).

This research was funded by NSERC Discovery Grant: RGPIN-2020-06235 to the Claycomb lab. Several trainees and staff who contributed to this work were funded by: CIHR Project Grants PJT-178076 and PJT-186154 to the Claycomb lab.

